# Leveraging epigenetic vulnerabilities of the stem cell-related *HOX*-signature in glioblastoma

**DOI:** 10.64898/2026.05.26.727851

**Authors:** Davide Chiesi, Pierre Bady, Mihalis V. Xirouchakis, Carla Mendes Ferreira, Kedir S. Mohammed, Monika E. Hegi

**Affiliations:** Neuroscience Research Center and Service of Neurosurgery, Lausanne University Hospital and University of Lausanne, Lausanne, Switzerland; L. Lundin and Family Brain Tumor Research Center, Departments of Oncology and Clinical Neurosciences, Lausanne University Hospital and University of Lausanne, Lausanne, Switzerland; Translational Data Science & Biomedical Data Science Center, Lausanne University Hospital and University of Lausanne, Lausanne, Switzerland; SIB Swiss Institute of Bioinformatics, Lausanne, Switzerland

**Keywords:** HOX-signature, super-enhancer, epigenetic vulnerability, drugability, glioblastoma

## Abstract

**Background:** Glioblastoma (GB) is the most aggressive primary brain tumor, characterized by therapy resistance, attributed to a multitude of epi-genetic changes resulting in phenotypic plasticity with altered cell states. To uncover druggable epigenetic vulnerabilities, we disturbed GB-derived spheres and observed coordinated repression of the aberrantly activated hemopoietic stem-like cell signature, dominated by *HOXA* genes. This signature has been associated with poor prognosis and resistance to therapy in GB. Here we investigate biological vulnerabilities associated with the deregulated epigenetic landscape in high-*HOX* GB.

**Methods:** GB-derived spheres (GS) were treated with an inhibitor of Bromodomain and extra-terminal motif proteins (BETi) (JQ1) or transduced with inducible constructs to genetically modulate *HOXA10* expression (shRNA for knockdown, ectopic *HOXA10*). Functional effects were evaluated through proliferation, neurosphere formation, and senescence assays. Epigenomic profiling incorporated RNA-seq, ChIP-seq, ATAC-seq, promoter capture MicroC, and DNA methylation.

**Results:** BETi-mediated rapid, coordinated downregulation of the *HOX-*signature, suggested direct transcriptional regulation. Knockdown of *HOXA10* alone yielded similar effects, decreasing expression of *HOXA* genes, reducing proliferation, self-renewal capacity, and triggering senescence. Conversely, ectopic *HOXA10* expression was ineffective in reactivating the *HOXA* cluster, or reverse BETi-mediated biological effects. Integrative epigenomic analysis of high-*HOX*-GS revealed concerted activation of the *HOXA* region, with broad domains of H3K27ac/H3K4me3 associated with super-enhancer activity, open chromatin (ATAC) and focal DNA hypomethylation. Architectural changes included altered CTCF interactions and increased promoter-anchored looping.

**Conclusion:** These results position the *HOX-*signature as a potential therapeutic target and offer a mechanistic rationale for disrupting BET-dependent transcriptional regulation in high-*HOX* GB.

**Key points:**

- Epigenetic activation of stem cell-related high-*HOX* signature in GB is associated with a super-enhancer encompassing the *HOXA* locus.
- Targeting this vulnerability by BETi or *HOXA10* knockdown results in concerted repression and loss of stemness features.

**Graphical Abstract:** 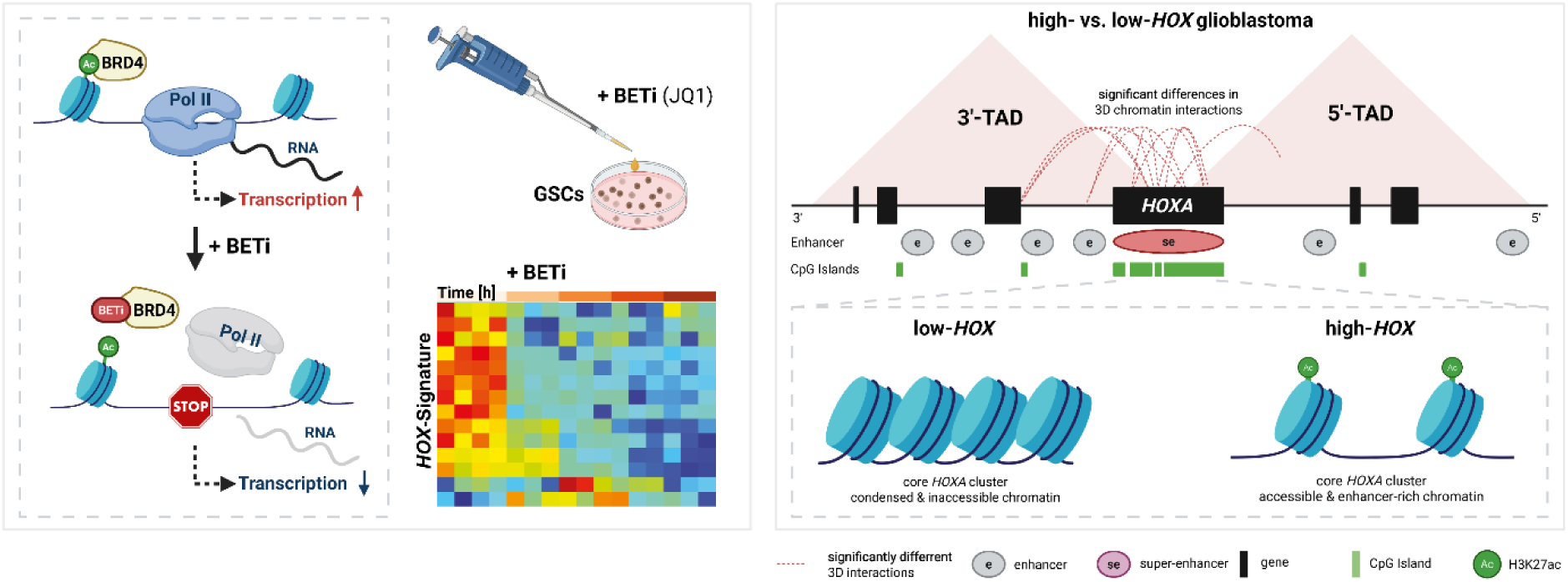

Created in BioRender. Chiesi, D. (2026) https://BioRender.com/eknk0ez

**Importance of study:** Glioblastoma (GB) are the most aggressive brain tumors in adults that are difficult to treat, due to their high plasticity resulting invariably to resistance to therapies. Here we report on the identification of epigenetic vulnerabilities that may be leveraged in combination therapies. Disturbing GB-derived stem-like cells with epigenetic drugs, we uncovered that a *HOXA* gene dominated hematopoietic stem cell-related signature, previously associated with aggressiveness and treatment resistance, can be repressed in a coordinated manner, resulting in loss of stem cell features. Analysis of the underlying epigenetic landscape revealed that the *HOXA* region was activated in high-*HOX* glioblastoma through the formation of a super-enhancer. This feature presents a particular vulnerability that may be leveraged by BETi as strategy of a combination therapy.

## 2. Introduction

Glioblastoma (GB) *IDH-*wildtype is the most common and most aggressive primary brain tumor combined chemoradiotherapy, and the addition of tumor treating fields ^1^. Efforts evaluating the addition of targeted therapies or various immunotherapies have failed so far. Treatment resistance has been attributed, at least in part, to the extensive genetic and epigenetic heterogeneity of GB, as well as the capacity of tumor cells to evade treatment through intrinsic or therapy-induced plasticity ^2^.

Aberrant epigenomic alterations play a central role in the pathogenesis of GB, contributing to transcriptional plasticity ^2,3^. In addition to cancer-associated genetic alterations, epigenetic changes characterize tumor subtypes, including cell of origin specific features, and phenotypic adaptations that contribute to treatment resistance, making them attractive targets for therapeutic intervention and biomarker development ^4^.

Pharmacological perturbation of epigenetic regulators has therefore emerged as a strategy not only to inhibit tumor growth, but also to expose transcriptional vulnerabilities that sustain malignant programs in GB and may serve as actionable targets.

Bromodomain and extra-terminal motif (BET) proteins constitute a family of epigenetic “reader” proteins that recognize acetylated lysine residues on histone tails and promote transcription by recruiting transcriptional co-activators and elongation machinery to active promoters and enhancers ^5^. BET proteins, in particular BRD4, play a key role in sustaining transcription at highly active loci and have been implicated in maintaining oncogenic transcriptional programs across multiple cancer types ^6^. BET inhibitors (BETi) have shown preclinical efficacy in GB models, with some compounds displaying favorable blood–brain barrier penetration and have been under clinical investigation in patients with glioblastoma ^7–9^. Beyond their antiproliferative effects, BETi have been shown to modulate specific transcriptional programs in GB models, including coordinated repression of interferon-stimulated gene signatures and pathways linked to treatment resistance, such as repression of *MGMT* in *MGMT* promoter unmethylated GB cells, highlighting their utility as functional probes of epigenetic vulnerabilities ^10,11^.

Among the transcriptional programs aberrantly activated in GB, dysregulated expression of *HOX* genes has been observed. These master regulators of embryonic patterning and cell fate are not expressed in the forebrain during development, nor in the adult brain. Coordinated aberrant expression of this hematopoietic stem cell-related signature ^12^ has been reported in subsets of GB and linked to poor prognosis, stemness, therapy resistance and recognized as a potential druggable target ^13–16^. Transcriptomic studies have identified coordinated *HOX* gene expression signatures in GB, suggesting that these loci may be regulated as a coordinated functional unit rather than as isolated genes ^14,17–19^. However, the epigenetic mechanisms that sustain aberrant *HOX* activation in GB and the extent to which these programs represent actionable transcriptional dependencies remain incompletely understood. Building on these observations, Kurscheid *et al.* demonstrated that coordinated *HOX* gene expression in GB was tightly linked to locus-specific DNA methylation patterns, with a central role for *HOXA10,* in the context of the GB-characteristic gain of chromosome 7 (CHR 7) ^17^. The aberrant epigenetic upregulation of the *HOX* genes, stratifies tumors into biologically and clinically distinct subgroups ^13–15,17,19^.

Here we investigated the functional and epigenetic features associated with the coordinated repression of the stem cell-related *HOX* signature observed in GB-derived sphere lines (GS) upon perturbation with the BET inhibitor JQ1. We identified the *HOXA-*dominated *HOX* signature transcriptional program as a reproducible and sensitive target in high-*HOX* GS lines using BETi or interfering with the *HOX* transcriptional program by knockdown of *HOXA10*. Integrating transcriptomic, epigenomic and functional analyses, we shed light on the epigenetic landscape underlying the coordinated activation and repression of *HOXA* in GS that may uncover druggable vulnerabilities.

## 3. Materials & Methods

### Cell culture

Patient-derived glioblastoma sphere (GS) lines LN-2540GS (RRID: CVCL_WS28), LN-2669GS (CVCL_WS29), and LN-2683GS (CVCL_WS30) were established, molecularly characterized and authenticated in our laboratory following institutional ethics guidelines (Brain Tumor Biobank BB 031_BBLBGT, CER-VD protocol F25/99); and cultured under stem cell conditions ^17,20,21^. HEK293T cells were maintained in DMEM with Glutamax and 5% FBS. All lines were regularly tested for mycoplasma. The BET inhibitor JQ1 (APExBIO, No. A1910) was used at the indicated concentrations. Additional details are available in the Supplementary Methods.

### Plasmid Preparation and Molecular Cloning

Tet-ON inducible *HOXA10-tGFP* expression constructs were made and transduced into the GS lines. In brief, the *HOXA10-tGFP* insert was PCR-amplified from pCMV6-AC-HOXA10-tGFP and cloned into a modified pCW22-Cas9-Blast vector through In-Fusion® Snap Assembly (Takara Bio Inc., 638948). Constructs were verified by sequencing. Primer sequences are listed in Table S1, and additional details can be found in the Supplementary Methods.

### Lentiviral Particle Production and Transduction

Inducible shRNA constructs targeting *HOXA10* (shHOXA10#2 and shHOXA10#5) and a non-targeting control (shNTC) were obtained from Horizon Discovery. Lentiviral particles were produced in HEK293T cells, concentrated, and used to transduce GS lines as described in the Supplementary Methods. Transduced cells were selected with puromycin (shRNA) or blasticidin (ectopic HOXA10-tGFP). Additional information on shRNA constructs is provided in Table S2.

### Proliferation Assay

Cell proliferation was assessed using the CyQUANT™ Direct Red Cell Proliferation Assay (Invitrogen™, C35013). Cells were seeded in a black, clear-bottom 96-well plate (Costar®, 3603) and treated the following day with DMSO, 0.5 µM JQ1, 0.5 µg/mL doxycycline, or combinations thereof. Fluorescence was measured on a SpectraMax® iD3 plate reader after a 1-hour incubation at 37°C and 5% CO₂. Readings were taken on days 0, 4, 8, 12, and 16, with the medium and treatment replenished on day 8. GS spheroids were centrifuged at 200 × g for 2 minutes before each reading.

### β-Galactosidase Activity Assay

Total β-galactosidase activity was measured as a senescence marker using the luminescent Beta-Glo® Assay System (Promega, E4720). Cell seeding and treatment conditions were identical to those in the proliferation assay and were performed in white 96-well plates (Costar®, 3917). Luminescence was measured on a SpectraMax® iD3 plate reader after a 30-min incubation at RT. Signal was normalized to relative cell number by dividing the β-galactosidase signal by the corresponding relative fluorescence value from the final time point of the proliferation assay.

### Neurosphere Formation Assay

The neurosphere formation assay was adapted from Gusyatiner *et al.* ^10^. For each biological replicate, technical triplicates of shNTC, sh*HOXA10*#2, and sh*HOXA10*#5 spheres were seeded into wells of a 24-well plate. The following day, cells were treated with DMSO, 0.5 µM JQ1, 0.5 µg/mL doxycycline, or a combination. Media and treatments were refreshed every 7 days. On day 21, spheres 50 µm in diameter were counted. The experiments were performed in biological triplicate.

### RNA Isolation and Real-Time Quantitative PCR (RT-qPCR) Analysis

Total RNA isolation and qRT-PCR were performed as previously described ^22^ using primers listed in Table S1. Expression levels were normalized to *GAPDH*.

### Western Blotting

GS lines were collected by centrifugation. Westerns were performed as previously described ^22^ and probed with primary antibodies and corresponding secondary antibodies, as detailed in Table S3.

### RNA-seq Sample and Library Preparation, Sequencing, and Preprocessing

RNA-seq data from LN-2683GS spheres treated with 1 µM JQ1 or DMSO over a time course of 48h were obtained from Gusyatiner *et al.* ^10^ (GEO accession number, GSE329546). For the transduced LN-2683GS (shNTC, sh*HOXA10*#2, and sh*HOXA10*#5 spheroids, RNA-seq was performed following doxycycline and JQ1 treatment. Libraries were prepared and sequenced at the Genomic Technology Facilities (GTF, University of Lausanne). Data were processed using the nf-core/rnaseq pipeline, aligned to GRCh38, and normalized with trimmed mean of M-values (TMM). Filtered genes (Table S4) were subjected to differential expression analysis using a generalized linear mixed model with a negative binomial distribution where construct type and replicate (batch) information are included as random intercepts. Genes with q-value ≤ 0.05 and |log2 fold-change| ≥ 0.58 were considered significant. Pathway over-representation analysis (ORA) was performed using MSigDB gene sets, Bonferroni-adjusted P-value ≤ 0.05 ^23,24^. Additional details on library preparation, sequencing, bioinformatic processing, statistical modeling, and pathway analysis are provided in the Supplementary Methods. (Acc# GSE329531)

### Chromatin Immunoprecipitation (ChIP)

ChIP-seq data and corresponding RNA-seq data for LN-2683GS spheres treated for 2h with JQ1 or DMSO were obtained from a previous study ^10^ (GEO: GSE329543, GSE329544). For ChIP-seq analysis of LN-2540GS and LN-2669GS spheroids in this study, chromatin was prepared using the ChIP-IT® Express Magnetic Chromatin Immunoprecipitation Kit (Active Motif, 53008) according to the manufacturer’s suspension cell protocol.

For ChIP-qPCR, 7 µg of chromatin per reaction was immunoprecipitated using antibodies listed in Table S3. Enrichment at target regions (primer sequences in Table S1) was calculated as a percentage over input and fold enrichment over IgG.

For ChIP-seq, 10 µg of chromatin was immunoprecipitated with antibodies detailed in Table S3. Libraries were prepared by the GTF (University of Lausanne) and sequenced on an AVITI™ Sequencer. ChIP-seq data were processed using the nf-core/chipseq pipeline (v2.0.0) ^25^, aligned to GRCh38 with Bowtie2, and peaks were called using MACS2. CTCF data were analyzed with the sevenC R package to compute correlation-based chromatin contacts ^26^. Detailed protocols for ChIP-qPCR, ChIP-seq library preparation, and bioinformatic analysis are available in the Supplementary Methods. (Acc #GSE329530)

### ATAC-seq Sample and Library Preparation

LN-2540GS and LN-2669GS spheroids (1 × 10⁶ cells each) were collected and treated with DNase I (StemCell Technologies, 07900) to remove potential apoptotic artifacts. Libraries were prepared using the ATAC-seq Kit (Active Motif, 53150) according to the manufacturer’s instructions. After tagmentation and PCR amplification, libraries were cleaned with AMPure XP beads (Beckman Coulter, A63880) using a 1.8X ratio. Tagmentation and library preparation efficiencies were then tested on a Fragment Analyzer. Sequencing was performed on an AVITI™ Sequencer (150-nt paired-end, 100 million reads/sample). Data were processed using the nf-core/atacseq pipeline (v2.1.2), aligned to GRCh38 with Bowtie2 (v2.4.4), and peaks were called with MACS2 (v2.2.7.1). (Acc# GSE329528)

### Promoter Capture Micro-C

LN-2540GS and LN-2669GS spheroids were processed by Cantata Bio (Scotts Valley, CA, USA) using the Dovetail® Micro-C (Dovetail, 20101E) and Human Pan Promoter Panel v1.0 (Dovetail, part no. CP3-PP-001) Kits, with sequencing to ∼150 million paired-end reads. Reads were aligned to GRCh38. Contact matrices were processed using the cooler Python package and R packages *HiCExperiment*, *HiContacts*, and *HiCcompare*. A/B compartments were predicted using constrained K-means in HiCDOC. TADs were identified with *SpectralTAD*. Differential analysis between the two sphere lines was performed using *multiHiCcompare*, with interactions tested by exact binomial tests. Detailed protocols and analysis parameters are provided in the Supplementary Methods. (Acc# GSE329532).

### Illumina Methylation Data

The DNA methylation datasets were previously published (Illumina human 450 k, GEO: GSE60274; ^17^). The datasets corresponding to LN-2540GS, and LN-2669GS were re-annotated for GRCh38 using Illumina EPICv1 manifest. Probes with a detection p-value >0.01 in >1% of samples were removed. Normalized beta-values were used for visualization ^27,28^.

### Statistical Analysis

All statistical analyses were performed using R (version 4.5.2) or GraphPad Prism 10. The data were analyzed with the statistical test described in the corresponding figure legend. Statistical significance was determined based on p-values, as indicated by asterisks in the figures: p ≤ 0.05 (*), p ≤ 0.01 (**), p ≤ 0.001 (***), and p ≤ 0.0001 (****). Data are presented as mean values, and error bars represent the standard deviation (SD).

## 4. Results

### Concerted downregulation of *HOXA* dominated stem-cell signature by the BET Inhibitor JQ1 in High-*HOX* Stem-like Glioblastoma-Derived Spheres

To identify epigenetic deregulation in GB, we analyzed RNA sequencing (RNA-seq) data from the patient-derived LN-2683GS sphere line treated with the BET inhibitor JQ1 over a 48-hour time course, as previously reported by Gusyatiner *et al.* ^10^. This analysis revealed dynamic transcriptional response patterns, of which two showed rapid and sustained downregulation of expression ^11^. Among these significantly repressed genes, we identified the stem cell-related *HOX* signature that was coordinately downregulated. This signature is enriched for *HOXA* genes, comprising, but not limited to, *HOXA3*, *HOXA5*, *HOXA7*, *HOXA9*, and *HOXA10*, together with *HOXC6*, *HOXD3*, *HOXD8*, *HOXD10*, and the stem cell marker *PROM1* (CD133) ^14^. Gene Set Enrichment Analysis (GSEA) against the glioblastoma-derived molecular signatures from Murat *et al.* ^14^ revealed a significant negative enrichment score for the *HOX* signature (G28) following JQ1 treatment (Figure S1).

To place these transcriptional effects into their epigenetic context, we investigated the epigenetic landscape in the *HOXA* region on CHR 7. The DNA methylation profile of LN-2683GS spheres closely matched that of the corresponding patient’s GB (GB 2683), emphasizing the clinical relevance of this *in vitro* model (Figure 1A). In untreated spheres, the *HOXA* cluster exhibited a permissive epigenetic configuration, characterized by focal DNA hypomethylation and broad enrichment of active histone marks (H3K27ac, H3K4me3), consistent with robust, concerted transcriptional activation (Figure 1B). BETi with JQ1 for 2 h reduced BRD4 binding and decreased binding of RNA Polymerase II (Pol2), associated with downregulation of gene expression across the entire *HOXA* cluster. These coordinated changes suggested that JQ1 treatment-induced rapid downregulation of *HOXA* cluster gene expression, indicating that the actively transcribed *HOXA* cluster genes are highly sensitive to BETi in this high-*HOX* GS line (Figure 1).

**Figure 1.**
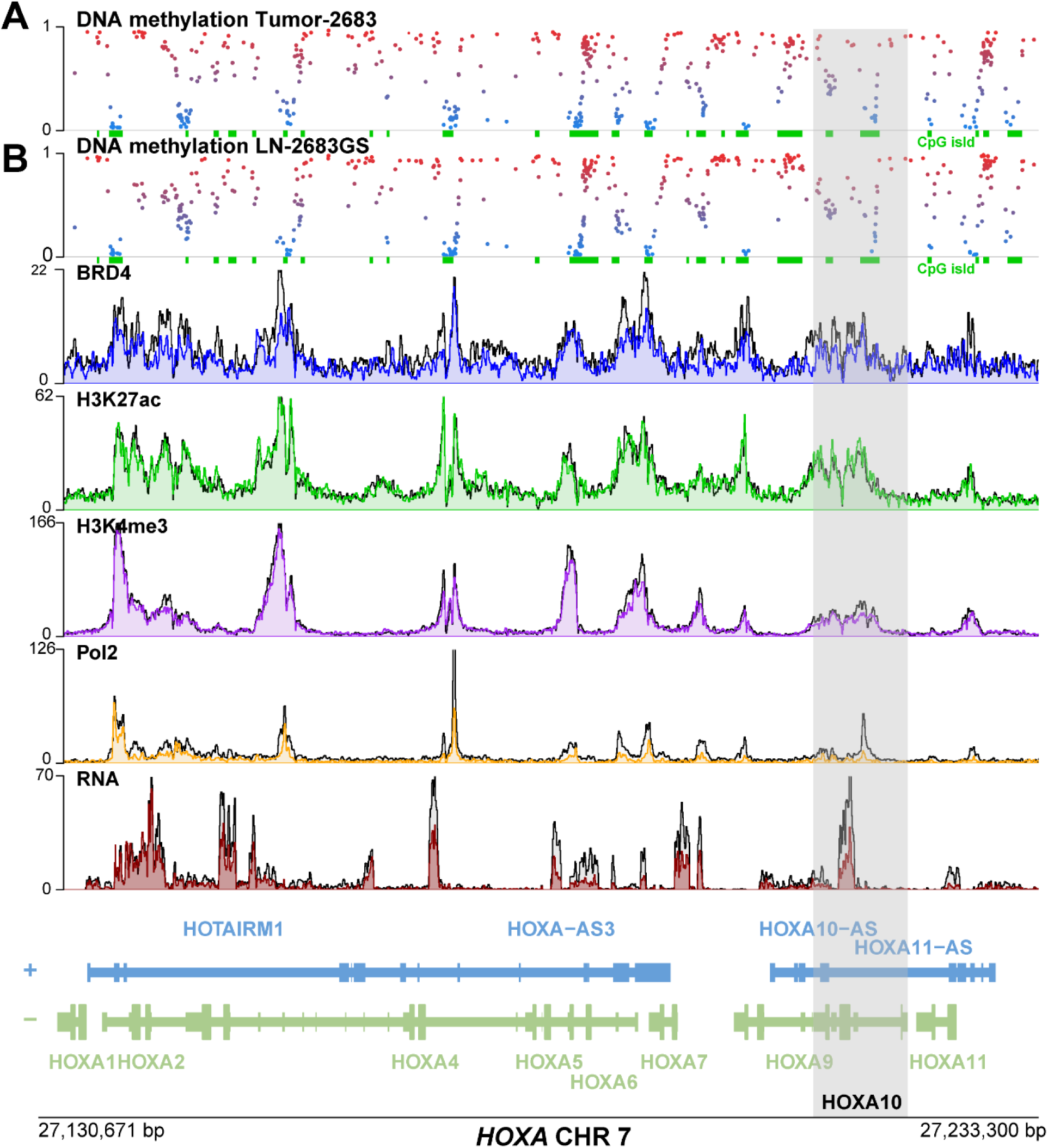
The concerted expression of *HOXA* genes is repressed by BETi in high-*HOX* glioblastoma-derived spheres. (A ) The DNA methylation profiles (HM-450K) in the *HOXA* cluster region on CHR 7 are shown for GB-2683 and the corresponding derived sphere line LN-2683GS. CpG Islands are indicated as green squares. ChIP-seq and RNA-seq data analyzed here, were extracted from a previously reported data set of LN-2683GS ^10^ that had been treated with 1µM JQ1 for 2 h, depicted in color, untreated control in black. **(B)** The coverage of ChIP-seq data is visualized for BRD4, H3K27ac, H3K4me3, RNAPII, and for the corresponding RNA-Seq data. Data shown correspond to the mean of three biological replicates. Profiles are shown for a bin size of 50 bp, alignment GRCh37.

The JQ1 sensitivity of high-*HOX* GS lines to coordinated downregulation of *HOXA* gene expression was confirmed in LN-2683GS and LN-2669GS. Accordingly, depletion of HOXA10 protein was observed over a 120 h time course (Figure S2).

### *HOXA10* knockdown is associated with the repression of other *HOXA* genes

Intrigued by the coordinated repression of *HOXA* gene cluster expression by BETi in high-*HOX* GS lines, we investigated whether targeting an individual gene within the locus would affect cluster-wide expression. We focused on *HOXA10*, whose promoter DNA methylation pattern in two GB cohorts showed one of the strongest correlations with the global *HOX* signature expression, as we reported previously ^17^. Doxycycline (DOX)-inducible shRNA-mediated knockdown of *HOXA10* in the high-*HOX* GS lines LN-2683GS and LN-2669GS efficiently reduced *HOXA10* mRNA and protein levels within 72 h and 120 h, respectively (Figure 2). Moreover, *HOXA10* knockdown led to coordinated downregulation of other *HOXA* cluster genes, including *HOXA5*, *HOXA6*, *HOXA7*, and *HOXA9*, which was reproduced in LN-2669GS. The effect of coordinated downregulation by the sh*HOXA10s* was comparable to that of JQ1 treatment. Notably, combining *HOXA10* knockdown with JQ1 treatment did not further reduce *HOXA* gene expression relative to either perturbation alone, as reflected also in similar depletion of the HOXA10 protein levels within 120 h.

**Figure 2.**
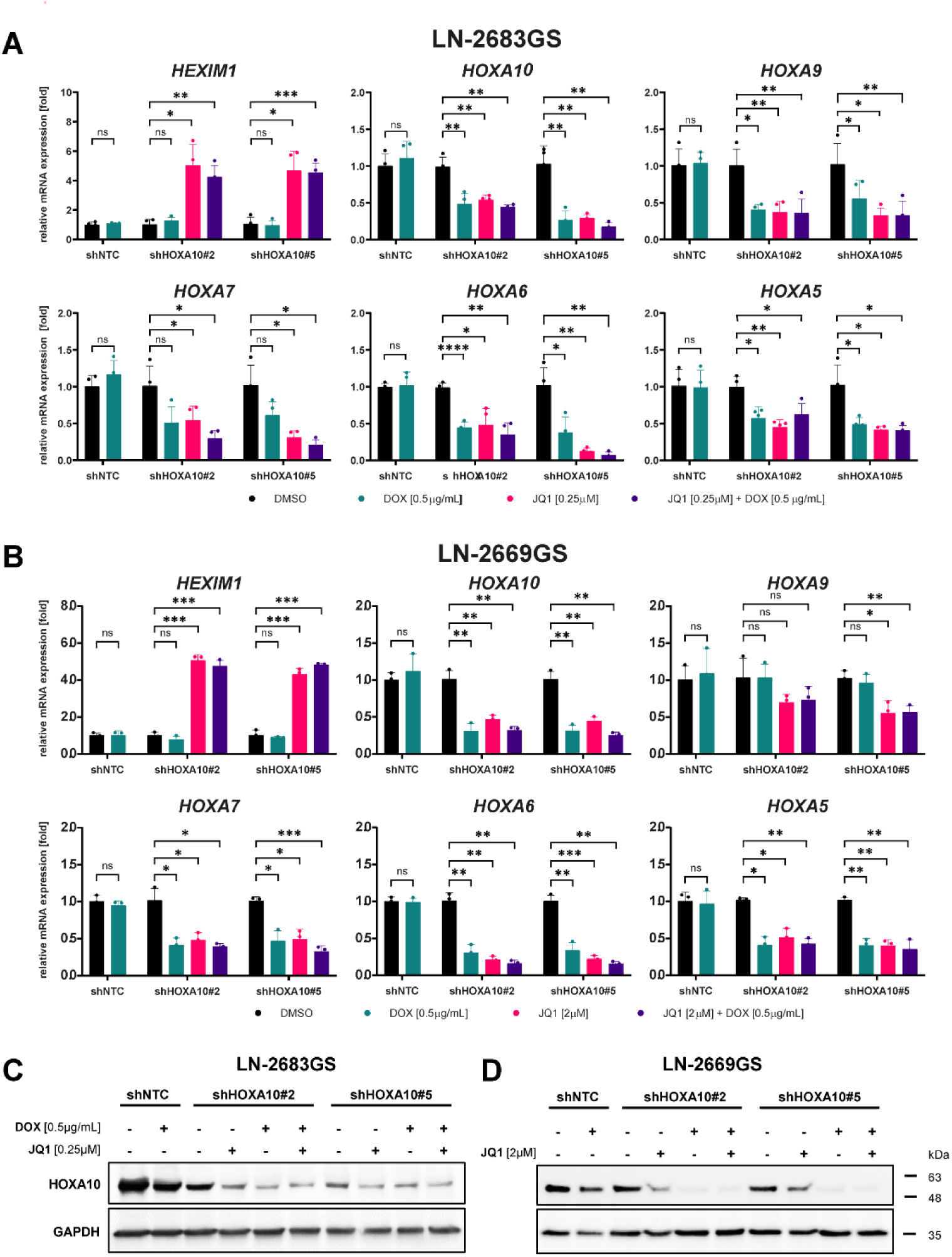
***HOXA10* knockdown is associated with reduced expression levels of other *HOXA* genes in high-*HOX* sphere lines LN-2683GS and LN-2669GS.** The two GS-lines transduced with the doxycycline-inducible *shHOXA10* (#2 or #5) or the control *shNTC* were treated with doxycycline and/or JQ1 at the concentrations indicated. Expression of *HOXA10* and other selected *HOXA* genes were determined after 72 h by qRT-PCR **(A, B)**. BET target engagement by JQ1 was monitored by measuring the induction of the pharmacodynamic marker *HEXIM1* ^51^. Results are shown as mean values of biological replicates (n=4 (A), n=3 (B), Error bars indicate SD). The data were normalized to the mean *GAPDH* expression and the corresponding control (DMSO). Treatment responses were tested by multiple unpaired t-tests with Welch’s correction. The protein levels of HOXA10 were measured after 120 h by Western, controlled by GAPDH **(C, D).** Representative biological replicates are shown (n=3). ns = non-significant, * = (p ≤ 0.05), ** = (p ≤ 0.01), *** = (p ≤ 0.001)

### *HOXA10* Knockdown Reduces Proliferation, Self-Renewal Capacity, and Induces a Senescence-Like Phenotype

Next, we determined the functional consequences of *HOXA10* downregulation in the high-*HOX* GS-lines. DOX-induced *HOXA10* knockdown significantly reduced cellular proliferation over 16 days in both LN-2683GS and LN-2669GS (Figure 3). Likewise, the neurosphere formation assays revealed reduction in secondary sphere formation, in the range of 50% after *HOXA10* knockdown, indicating a substantial loss of self-renewal capacity. The reduction of these biological features associated with malignancy was accompanied by induction of a senescence-like phenotype, evidenced by a 4- and 2-fold increase in β-galactosidase activity after 16 days in LN-2683GS and LN-2669GS, respectively. In accordance, decrease of the senescence-associated biomarker Lamin B1 (*LMNB1)* was observed at both the mRNA and protein levels within three and five days of *HOXA10* knockdown, respectively. These phenotypic changes mirrored those induced by BETi (JQ1).

**Figure 3.**
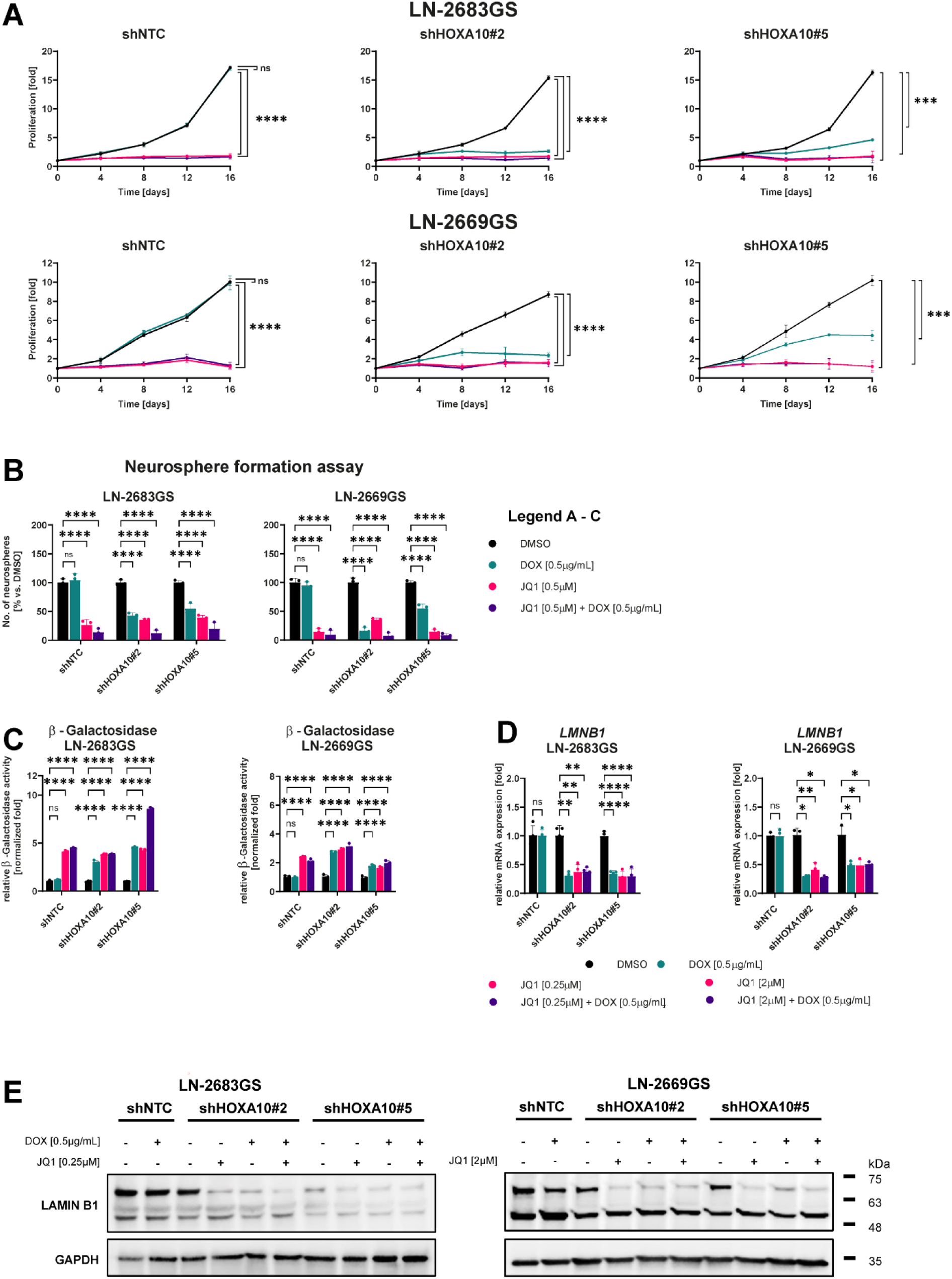
Functional effects of *HOXA10* knockdown and JQ1 treatment on high-*HOX* LN-2683GS and LN-2669GS spheres. The two high-*HOX* GS-lines, LN-2683GS and LN-2669GS, transduced with the inducible sh*HOXA10* #2, #5, or the non-targeting shNTC, were induced by doxycycline and/or treated with JQ1 at the concentrations indicated: **(A)** Relative proliferation of LN-2683GS or LN-2669GS was measured over 16 days. Treatment responses on day 16 were tested and expressed as fold changes relative to day 0. Results are shown as mean values of biological replicates (n = 3). Error bars indicate SD. Two-way ANOVA was tested, and Dunnett’s multiple comparisons test was used for post hoc analysis. **(B)** Neurosphere formation was assessed at 21 days of treatment (same as in **A**) for both high-*HOX* GS lines. Spheres ≥ 50µm were counted and normalized to the control (DMSO). **(C)** The senescence-like phenotype was investigated by measuring the relative β-Galactosidase activity after 16 days of treatment (as in A). Luminescence was normalized to the relative number of cells. **(D)** *LMNB1* expression was measured by RT-qPCR after 3 days of treatment as indicated. The data were normalized to the mean *GAPDH* expression and the corresponding control (DMSO). Treatment responses were tested by multiple unpaired t-tests with Welch’s correction. **(E)** Western blot analysis shows Lamin B1 protein expression after 5 days of treatment, as in (D), controlled by GAPDH. Representative biological replicates are shown (n = 3). ns = non-significant, * = (p ≤ 0.05), ** = (p ≤ 0.01), *** = (p ≤ 0.001), **** = (p ≤ 0.0001).

### Ectopic *HOXA10* expression is insufficient to activate the *HOX* transcriptional program or counteract BET inhibition

To assess whether *HOXA10* is sufficient to drive expression of the *HOX* gene signature, we introduced DOX-inducible ectopic *HOXA10-tGFP* expression into low- and high-*HOX* GS-lines. Ectopic expression of *HOXA10* had no effect on transcription of other *HOXA* genes, in the low-*HOX* GS-line LN-2540GS (Figure S3) nor in the high-*HOX* GS-lines LN-2669GS and LN-2683GS that are “wired” to express *HOX* genes (Figure S4). Moreover, the ectopic expression of *HOXA10* in the two high-HOX GS-lines was neither able to rescue cell growth, nor to prevent induction of the senescence-like phenotype upon JQ1 treatment. These findings indicate that ectopic *HOXA10* expression alone is insufficient to re-establish the *HOX* signature transcriptional program or to counteract BETi-induced effects.

### Transcriptomic Profiling after HOXA10 Depletion Reveals Repression of Stemness and Induction of Neural-Like Transcriptional Signatures

To characterize the transcriptional consequences of HOXA10 depletion, we performed RNA sequencing on LN-2683GS spheres five days after induction of sh*HOXA10*, or after five days of JQ1 treatment, when depletion of the HOXA10 protein is observed (Figure 2).

Analysis of the JQ1 response revealed that prolonged BETI induced widespread transcriptional reprogramming, confirming consistent transcriptional changes in key pathways after 5 days, extending beyond the previously described observation interval ^10,11^. Expression changes following the knockdown by *shHOXA10* at the time of HOXA10 depletion were more limited. Differential expression analysis identified 233 significantly altered genes (DEGs; 194 upregulated and 39 downregulated using a generalized linear mixed model (Bonferroni, p ≤ 0.05; |logFC| > 0.58) (Table S4). Overrepresentation analysis (ORA) of cancer-related signatures (MSigDB H and C2) revealed enrichment of pathways associated with stemness, differentiation, and ECM organization – all of which overlapped with JQ1-affected gene sets already present after 48 h of treatment. Interestingly, top pathways identified by ORA using Gene Ontology signatures (C5:GO_BP and C5:GO_CC) showed significant enrichment for neuronal differentiation, axonogenesis, and synaptic signaling programs (Figure S5; Table S5). We compiled a prioritized shortlist of candidate genes from the RNA-seq data for validation. Selection was based on fold-change (logFC) in significant DEGs, their consistent contribution to enriching stemness and neuronal differentiation signatures, and their baseline expression levels in the GS-lines. This yielded a panel of genes with established roles in neural development or GB biology, including downregulation of the stemness-associated gene *Distal-Less Homeobox 1* (*DLX1)* ^29^, alongside induction of genes with established neurogenic functions. including the transcription factor *Neuronal Differentiation 1 (NEUROD1)* ^30,31^, the tumor suppressor *KLF Transcription Factor 9* (*KLF9*) ^32,33^, the neuronal markers *L1 Cell Adhesion Molecule* (*L1CAM)* ^34,35^ and the *Synaptosome Associated Protein 25 (SNAP25)* ^36^ (Figure S5). These genes were absent from those differentially expressed after short-course JQ1 exposure, indicating that they likely correspond to secondary effects of *HOXA10* loss that are not captured as early transcriptional responses to BETi.

### Local organization of chromatin architecture, histone modifications at regulatory regions of the *HOXA* locus in high-*HOX* spheres

To explore whether the transcriptional activation of the *HOXA* cluster in high-*HOX* GB is linked to differences in chromatin structure, we characterized and compared the epigenetic landscape in the low-*HOX* LN-2540GS and high-*HOX* LN-2669GS GS lines at several hierarchical levels of chromatin organization.

The three-dimensional chromatin structure was investigated using promoter capture Micro-C. Compartmentalization of CHR 7 into A/B, assigned the *HOXA* locus to the B compartment (at 516 kb resolution) in both LN-2540GS and LN-2669GS, like most of the distal part of CHR 7p, including the *EGFR* locus (Figure S6A). Of note, both GS lines display classic GB hallmarks of gain of CHR 7 and loss of CHR 10, based on 450 k data. No major differences were apparent in higher-order chromatin organization. Differential analysis of significant local interactions on CHR7 revealed a peak at the *HOXA* locus and at the *EGFR* locus, with opposite signs in the two GS-lines. This is in accordance with the known amplified *EGFR* locus of the low-*HOX* GS-line LN2540GS (Figure S6B).

At the level of TADs, at 64 kb and 32 kb resolution, the *HOXA* locus was located at the intersection of two TADs. Local interactions within the *HOXA* region were increased in the high-*HOX* LN-2669GS (Figure 4A). Furthermore, these areas, spanning the *HOXA1*-*HOXA3* and *HOXA9*-*HOXA10* regions, were associated with enhanced chromatin accessibility in the high-*HOX* GS line, based on ATAC-seq coverage (Figure 4B). The coverage of CTCF ChIP-seq identified some differences in detected CTCF peaks at the 5’ end of the *HOXA* region (#8) or outside (#9) in the high-*HOX* LN-2669GS that were absent in the low-*HOX* GS-line or vice versa, suggestive of distinct, unique loops linking these CTCF peaks (#8, #9) to upstream *HOXA* regions (3’). LN-2669GS showed unique ATAC-seq enrichment across multiple regions of the *HOXA* cluster, concordant with the regions corresponding to sites of increased chromatin interactions identified by Micro-C. The significant differences (fold changes) of these interactions between the GS lines are illustrated for the extended *HOXA* region in Fig. 5 (locations summarized in Table S6). Differences also appeared for contacts linking *HOXA* promoters to *SKAP2* (250–350 kb) at the 3′ TAD boundary of the *HOXA* region that is part of the GB-associated *HOX* signature. In concordance, these regions with increased interactions exhibited open chromatin (ATAC) and enrichment of H3K27ac_and H3K4me3 coverage, suggesting activated regulatory elements across the *HOXA* cluster in the high-*HOX* spheres. Moreover, enhancer classification suggested that a large part of the *HOXA* cluster, from *HOXA1* to *HOXA9*, was associated with super-enhancer activity, along with sharp promoter-proximal peaks for H3K27ac and H3K4me3. The corresponding DNA CpG methylation patterns of the two GS lines aligned with the respective activated or silenced state of the *HOXA* regions.

**Figure 4.**
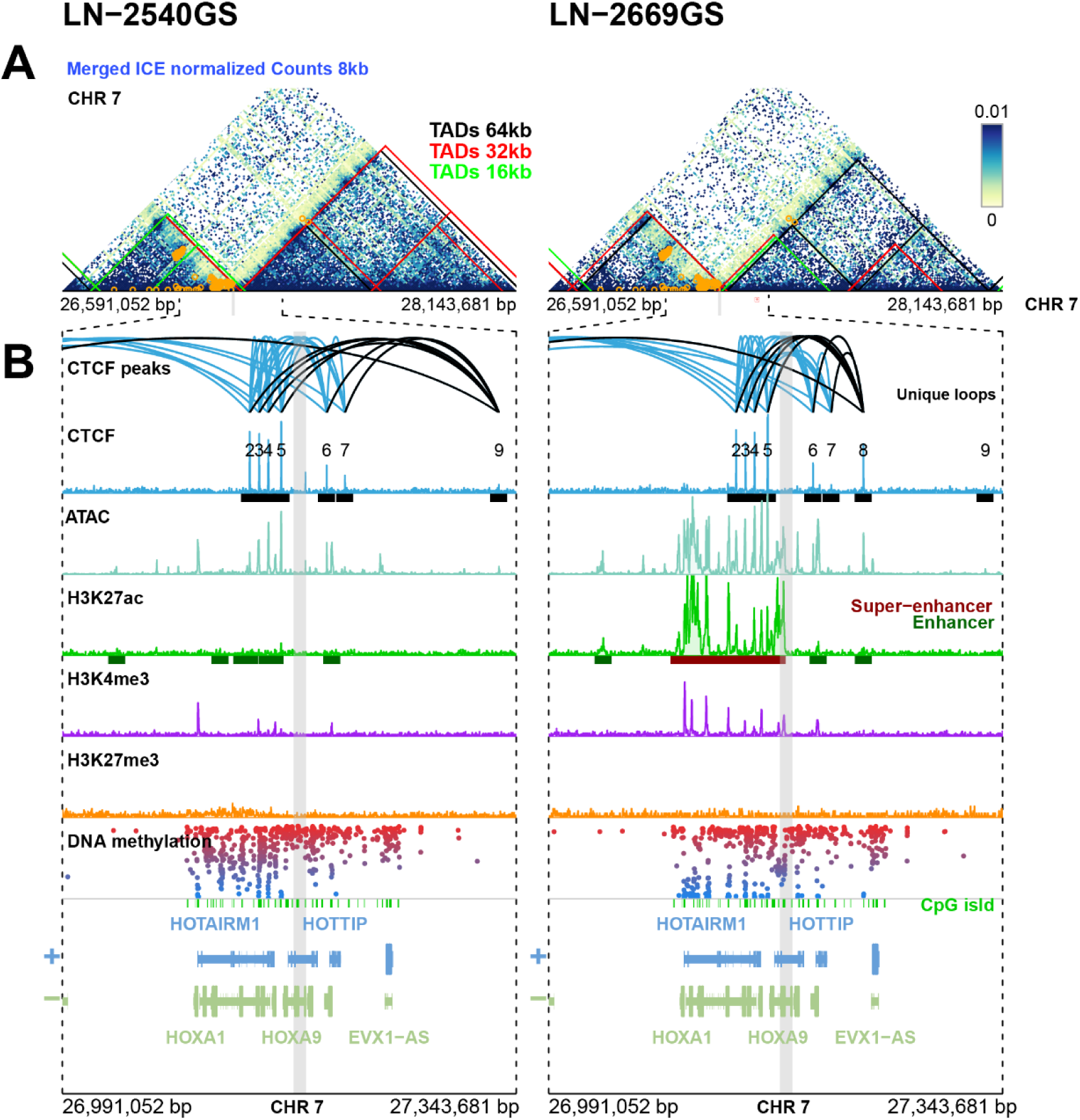
Comparison of epigenetic landscapes in the *HOXA* region between the low-*HOX* LN-2540GS and high-*HOX* LN-2669GS sphere lines. LN-2540GS and LN-2669GS are represented in blue and orange, respectively. **(A)** Three-dimensional interactions from Promoter Capture Micro-C data are visualized as a heatmap at 8-kb resolution (ICE-normalized counts [iterative correction for the HiC matrix], averaged by GS line, 3 replicates each). TADs obtained by *spectralTAD* are superimposed on the heatmap at 16 kb (green), 32 kb (red), and 64 kb (black) resolution. Significant genomic interactions from the differential comparison (exact binomial test) between both GS lines are marked with orange circles (abs(logFC) > 1, p-value adjusted by Bonferroni correction < 0.1) (detailed locations and statistics in Table S6). **(B)** CTCF peaks detected by peakcaller MACS2 are represented as black rectangles (numbered). Unique CTCF loops are indicated in black. ATAC-seq coverage profiles indicate chromatin accessibility. Corresponding coverage for H3K27ac is shown, annotated for super-enhancers (dark red) and enhancers (green), respectively, as predicted by the ROSE algorithm. ChIP-seq coverage for H3K4me3 and H3K27me3. DNA methylation profiles (beta-values, 450k) for LN-2540GS and LN-2669GS are visualized with a color gradient, and corresponding CpG islands are indicated by green ticks. Alignment was performed for the Human Genome Assemblage version GRCh38. The grey shaded area indicates the position of the *HOXA10* gene.

**Figure 5.**
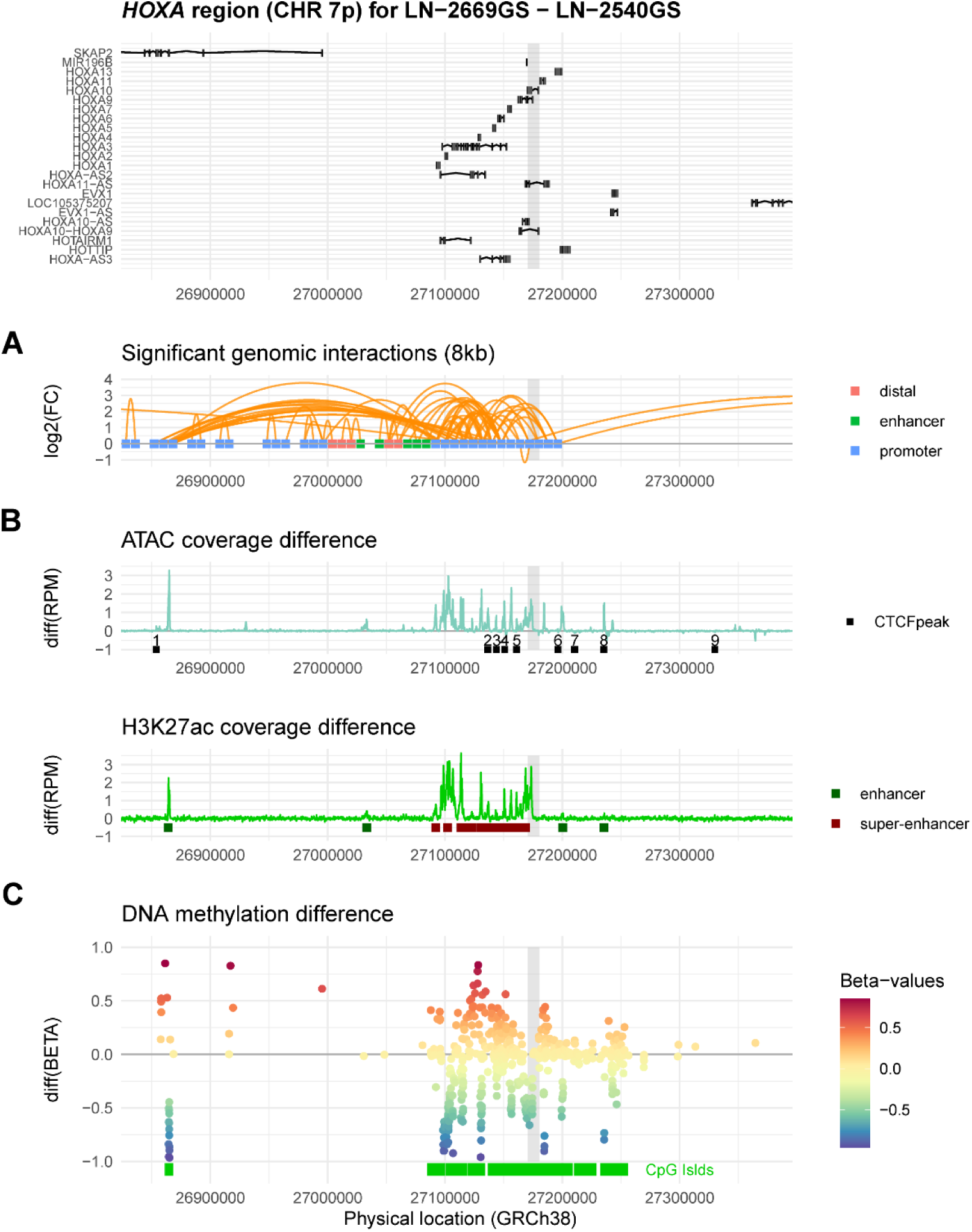
Differential genomic interactions in the extended *HOXA* region. **(A)** Significant genomic interactions from the differential comparison (exact binomial test) between the two GS lines, high-*HOX* LN-2669GS versus low-*HOX* LN-2540GS, determined by micro-C, are marked with orange arches (Suppl Table S6, detailed locations and statistics). The summit corresponds to log fold-change (FC; values > 0, correspond to higher interactions in the high-*HOX* LN-2669GS). The annotation for each probe is given by a color code for promoter (blue), enhancer (green), or distal (pink). **(B)** The difference of reads per million mapped reads (RPM) is provided for ATAC and H3K27ac coverage as a function of their physical location. The detected CTCF peaks and the enhancer classification by ROSE are superimposed on ATAC and H3K27ac tracks, respectively (enhancer, green, super-enhancer red). **(C)** The DNA methylation difference between the two sphere lines was computed with beta-values after functional normalization. CpG island locations are superimposed (green) on this track. The grey shaded area indicates the location of the *HOXA10* gene.

### Epigenetic landscape of other *HOX* clusters contributing to the high-*HOX* signature

To set the dominating contribution of the *HOXA* cluster into context of the overall *HOX* signature, the local epigenetic landscapes were visualized for the other *HOX* clusters (*HOXC*, CHR 12q; *HOXD,* CHR 2q; *HOXB*, CHR 17q). They contribute to varied degrees to the hematopoietic stem-like expression signature of high-*HOX* GB. Differential analyses of the high-*HOX* LN-2669GS versus the low-*HOX* LN-2540GS revealed the highest activity in the *HOXC* cluster, with concordant open chromatin (ATAC), active marks (H3K27ac) classifying as super-enhancers, and focal CpG hypomethylation. Some limited activity was observed in the *HOXD* cluster, without super-enhancer formation. Basically, no activity was observed in the *HOXB* cluster that does not contribute to the *HOX* signature. These observations are in line with their distinct contributions to the expression of the *HOX* signature (Figure S7A (*HOXC*); B (*HOXD)*; C (*HOXB*)).

### Activation of the canonical and alternative *HOXA10* promoter in the high-*HOX* GS line

Based on our previous findings that attributed a special role of *HOXA10* in the activation of the overall *HOX* signature in GB, we investigated the canonical and alternative *HOXA10* promoters in LN-2540GS and LN-2669GS by ChIP-qPCR (Figure S8). Enrichment of active histone marks (H3K27ac, H3K4me3, and H3K9ac) was significantly higher at both promoter regions in the high-*HOX* LN-2669GS than in LN-2540GS, with more pronounced differences (FC) at the canonical promoter. In accordance, enrichment of the repressive mark H3K27me3 was significantly lower in LN-2669GS across most promoter regions examined. These data confirm, at single-promoter resolution, that both the canonical and alternative *HOXA10* promoters are epigenetically active in the high-*HOX* GS line.

## 5. Discussion

Aberrant activation of a hemopoietic stem cell-like transcriptional program, dominated by *HOXA* genes, is a characteristic feature of a subset of GB. In this research, we reveal the epigenetic landscape in the *HOXA* region associated with this coordinated, activated transcriptional program in the high-*HOX* GB-derived sphere models. Functional perturbation experiments using pharmacologic interference with the epigenetically regulated transcriptional machinery via BETi revealed coordinated repression of the entire *HOX* signature. This was associated with reduced BRD4 occupancy and reduced Pol II binding. Profound effects were observed in the biology of high-*HOX* GS lines, reducing proliferation and self-renewal capacity and inducing a senescence-like phenotype. Most interestingly, similar biological changes were observed when the transcriptional program was genetically perturbed by targeting *HOXA10* alone.

Investigating the *HOXA* cluster with multi-layered epigenomic profiling of high- and low-*HOX* GS-lines revealed for high-*HOX* a permissive chromatin landscape with distinct CTCF loops, and increased chromatin interactions within the extended *HOXA* region. Alterations in CTCF binding have been linked to cancer-related enhancer rewiring ^37–39^. However, at a higher level of 3D organization of the chromatin, no major differences were observed.

The *HOXA* cluster is a gene and CpG island-rich region and forms a super-enhancer in the high-*HOX* GS-line (spanning *HOXA*1 to *HOXA*9) (Suppl Fig. S5). Interestingly, the active *HOXA10* is at the 5’ end of this super-enhancer. A special role has been attributed to *HOXA10,* as hypomethylation at its promoter showed the highest correlation with expression of the high-*HOX* signature in GB ^17^. Interference with *HOXA10* expression by shRNAs, was sufficient to repress cluster-wide activity of other *HOXA* genes, and phenocopied JQ1-mediated biological effects, reducing proliferation, self-renewal capacity, and induction of a senescence-like phenotype. However, ectopic expression was neither effective in activating other *HOXA* genes in low-*HOX* GS nor in rescuing high-*HOX* GS cells from JQ1-mediated biological effects. This suggests that the regulatory context of endogenous *HOXA10* expression is mechanistically important for the activity of the *HOX* signature. The regulatory context governing the coordinated regulation of *HOX* genes across the four distinct *HOX* clusters (A, C, D, B) located on different CHR remains unclear. However, their variable contribution to the overall *HOX* signature, dominated by *HOXA* genes, was well aligned with their local epigenetic landscapes, explaining the distinct activation levels and respective transcripts of the signature (genes, non-coding RNAs, microRNAs). These observations further emphasize the dominant role of the *HOXA* locus, which is associated with a super-enhancer that encompasses the entire region of the expressed *HOXA* genes.

Cluster-wide regulation is a defining feature of *HOX* loci during development outside the forebrain, where shared regulatory elements, chromatin looping, and coordinated epigenetic remodeling enable synchronized gene expression ^40^. Although *HOX* genes are not expressed in the development of the forebrain or the adult brain, the adoption of this aberrant concerted activation of the hemopoietic stem cell-related *HOX* signature in GB highlights the stem-like tumor state ^12,14,17,18^, in line with the reported loss of the repressive mark H3K27me3 across the four *HOX* clusters ^19^.

BETi rapidly suppressed cluster-wide expression, a change associated with decreased RNA Polymerase II occupancy, without concurrent alterations in histone modifications associated with active chromatin. This aligns with previous observations that BETi disrupts transcription, while enhancer-promoter contacts remain ^41^. This dissociation indicates that BET proteins are not necessary to establish the potential for transcription – unlike the open chromatin and enhancer landscape – but are essential for maintaining transcription at active loci ^5,6,42^. Furthermore, BRD4 binding is essential for the super-enhancer function, which can activate transcription of associated proto-oncogenes and maintain oncogenic identity in cancer ^43^. Super-enhancers have also been associated with the formation of loops, interchromosomal interactions, and higher-order hubs ^44^. The dependence on this continuous activation presents a potentially druggable vulnerability.

Genetic interference with shRNAs targeting *HOXA10* expression, a key component of the *HOXA* cluster, recapitulated the effects of BETi with similar functional outcomes and shared transcriptional programs. In particular, *HOXA10* knockdown affected pathways regulating stemness, differentiation, and ECM organization, as well as neuronal differentiation markers such as *NEUROD1*, *KLF9*, and *L1CAM*.

A key insight from this work is that the permissive landscape in high-*HOX* cells is defined by “local” coordinated epigenetic organization, comprising focal DNA hypomethylation, broad H3K27ac and H3K4me3 domains, and increased chromatin accessibility, associated with increased promoter-centered looping, especially in the *HOXA1-HOXA3* and *HOXA9-HOXA10* regions. In contrast, at a higher structural level, most of the distal part of CHR 7p, encompassing the *HOXA* cluster and the *EGFR* gene, was assigned to the B compartment that is associated with condensed chromatin (heterochromatin) and considered transcriptionally inactive ^45^. No difference in the compartmentalization of this genomic region was observed between the high- and low-*HOX* GS-line. Current efforts aim to identify higher-order structural epigenetic alterations (3D), including compartment switching that may give rise to new interactions to uncover druggable vulnerabilities in GB ^46,47^. Yet, local rewiring without compartmental switching has been recognized as a mechanism for inactivating repression of proto-oncogenes in cancer ^48,49^.

These findings identify targetable vulnerabilities and carry important therapeutic implications. They support the use of BETi to disrupt the *HOX*-driven stem-like state, while also complementing their known roles in repressing other programs relevant to GB ^7,50^, including interferon signaling ^10^ and MGMT expression ^11^. The similarities observed in phenotypes induced by *HOXA10* knockdown and BETi underscore the *HOXA* cluster as a vulnerability within a broader BET-dependent regulatory context in high-*HOX* GB. This indicates that *HOX* signature expression could be a useful biomarker for patient stratification in future epigenetic therapy trials. Our data provide a mechanistic rationale for strategies such as BETi or other epigenetic drugs that uncover vulnerabilities by destabilizing the epigenetic framework, thereby abrogating the stem-like state and possibly rendering cells more susceptible to differentiation or synergistic combination therapies.

## Acknowledgments

We thank the collaborators at the Genomic Technology Facility (GTF) of the University of Lausanne for their excellent technical support.

## Authorship statement

Conception and design: M.E.H., D.C., P.B.; Acquisition of data (experimental data) (D.C., M.V.X., C.M.F, K.S.M).; Analysis and interpretation of data D.C., P.B., M.E.H, M.V.X., C.M.F, K.S.M.; Computational biology and biostatistics, P.B.; Manuscript writing: D.C., M.E.H., P.B.; Review, and/or revision of the manuscript: all coauthors. Study supervision: M.E.H.

## Data availability

The datasets are available at the Gene Expression Omnibus (ncbi.nlm.nih.gov/geo/)

**Datasets, this study:** RNA-seq (LN-2683GS, transduced) GSE329531; ChIP-seq (LN-2669GS, LN-2540GS) GSE329530; ATAC-seq (LN-2669GS, LN-2540GS) GSE329528; microC (LN-2669GS, LN-2540GS) GSE329532.

**Previously described datasets:** Hu 450 k methylation data, GEO60274 ^17^; RNA-seq (LN-2683GS, BETi time course) GSE329546 ^10,11^; RNA-seq (LN-2683GS; associated with ChIP-seq) GSE329544 ^10^; ChIP-seq LN-2683GS; GSE329543 ^10^.

