## Supplemental Material &Tables for "Leveraging epigenetic vulnerabilities of the stem cell-related *HOX*-signature in glioblastoma": 20260522_HOX paper supplement_Submission.pdf

### 1. Supplementary Methods

#### Cell Culture

Patient-derived GB sphere (GS) lines LN-2540GS (RRID: CVCL_WS29), LN-2669GS (RRID: CVCL_WS29), and LN-2683GS (RRID: CVCL_WS30) were established and molecularly characterized in our laboratory [1-3] following institutional guidelines and approved by the Ethics Committee of the Canton de Vaud (CER-VD, protocol F25/99). All lines were regularly tested for mycoplasma contamination using the MycoAlert Kit (Lonza, LT07-418) and authenticated in 2025 by STR fingerprinting at the Forensic Genetics Unit of the University Center of Legal Medicine, Lausanne and Geneva [3]. GS lines were cultured under neural stem-like conditions in Dulbecco's Modified Eagle Medium/F12 (DMEM/F12; Life Technologies, 31331-028), supplemented with B27 (1X; Invitrogen™, 17504-001), 20 ng/mL epidermal growth factor (EGF; PeproTech, AF-100-15-100), and 20 ng/mL fibroblast growth factor (FGF; PeproTech, AF-100-18B-100). Adherent lines were cultured in DMEM with GlutaMAX (Gibco, 61965-026), supplemented with 5% fetal bovine serum (FBS; BioWest, S181B-500). Transduced cell or sphere lines were continuously maintained under the respective selection. The BET inhibitor JQ1 was dissolved in DMSO at 10 mM and added to cells at the indicated concentrations (APExBIO, A1910).

#### Plasmid Preparation and Molecular Cloning

Sphere lines LN-2540GS_HOXA10-tGFP$^{IND}$, LN-2669GS_HOXA10-tGFP$^{IND}$, and LN-2683GS_HOXA10-tGFP$^{IND}$ were created from parental lines using a Tet-ON system for ectopic *HOXA10-tGFP* expression. The pCW22-Cas9-Blast vector, provided by Prof. Joachim Lingner (EPFL) [4], was digested with SalI-HF (NEB, R3138S) and SbfI-HF (NEB, R3138S) to remove the Cas9 gene. The *HOXA10-tGFP* insert was PCR-amplified from pCMV6-AC-HOXA10-tGFP, sourced from OriGene (RG202939), and assembled with the linearized backbone using In-Fusion® Snap Assembly (Takara Bio Inc., 638948) (100 ng of total DNA per reaction, with a 2:1 ratio). The mixture was transformed into Stellar Competent Cells (Takara Bio Inc., 636764), and colonies with positive inserts were grown in Luria-Bertani (LB) broth containing 100 µg/mL ampicillin (Gibco™, 11593027). Plasmids were purified using the QIAGEN Plasmid Maxi Kit (QIAGEN, 12163) and confirmed by sequencing. Primer sequences are detailed in Supplementary Table S1.

#### Lentiviral Particle Production and Transduction

The SMARTvector Inducible Non-targeting Control (shNTC) and SMARTvector Human Inducible Lentiviral shRNA (shHOXA10#2 and shHOXA10#5) constructs were purchased as *E. coli* glycerol stocks from Horizon Discovery Ltd (shNTC: VSC11653; shHOXA10#2: clone ID V3IHSHEG_4915981; shHOXA10#5: clone ID V3IHSHEG_7259938). More information about shRNA constructs is available in Table S2.

GS cells were transduced using a protocol based on the Dharmacon™ Trans-Lentiviral shRNA Packaging Kit (Horizon Discovery, TLP5913), as previously described [5] and Frenster *et al.* [6]. Briefly, three 10cm dishes of HEK293T cells were first transfected with each construct using the kit's calcium phosphate reagent. Viral supernatant was collected at 24 and 48 h, filtered through a 0.45 µm Millex®-HA Filter Unit (Merck), and concentrated using Lenti-X™ Concentrator (Takara Bio, 631231). The concentrated virus was resuspended in media containing 8 µg/mL protamine sulfate (Sigma-Aldrich, P4505). Target GS cells were then incubated with the lentiviral suspension for 72 h. Selection was initiated 72 h post-transduction with puromycin (for shRNAs) or blasticidin (for ectopic HOXA10-tGFP), with the antibiotic concentration gradually increased to 10 µg/mL every 5 days.

#### RNA Library Preparation and Sequencing

$5 \times 10^5$ LN-2683GS_shNTC, LN-2683GS_shHOXA10#2, and LN-2683GS_shHOXA10#5 spheroids were seeded in 10 cm petri dishes. After 48 h, they were treated with 0.5 µg/mL doxycycline or control media. To confirm the efficient downregulation of HOXA10 protein induced by doxycycline in the shHOXA10 models, control Western blots were conducted in parallel for each biological replicate using $1 \times 10^6$ spheroids and the same treatment schedule. Total RNA was extracted using the ReliaPrep™ RNA Cell Miniprep System (Promega, Z6011), and ribosomal RNA was depleted prior to library preparation using the QIAseq FastSelect-rRNA HMR Kit (QIAGEN). rRNA-depleted RNA libraries were prepared and sequenced by the Genomic Technology Facilities (GTF, University of Lausanne).

#### RNA-seq Preprocessing and Normalization

RNA-seq data preprocessing was performed following the standard nf-core/rnaseq pipeline (version 3.11.2; https://nf-co.re/rnaseq/3.11.2/). After quality control, reads were aligned to the human reference genome (GRCh38, Ensembl release 113; hg38) using the HISAT2 aligner (version 2.2.1) and count data were provided by pseudo-alignment using Salmon (version 1.10.1).

Gene-level expression was summarized as trimmed mean of M-values (TMM)–normalized counts using the edgeR R package. Normalization included log-transformation and accounted for read counts and full library size [7,8].

**Differential Gene Expression Analysis**

The association between treatment and gene expression was assessed using a generalized linear mixed model (GLMM) based on a negative binomial distribution with log link. The model was defined as:

$$count \sim DOX + JQ1 + (1|type) + (1|replicate)$$

where DOX and JQ1 represent fixed effects for treatment conditions, and type (cell line) and replicate represent random intercepts that account for biological variability.

Differential expression analysis and statistical inference were performed using the glmmSeq R package (version 0.5.4; https://github.com/KatrionaGoldmann/glmmSeq), which models count-based RNA-seq data with random effects. Genes with q-values ≤ 0.05 and absolute log2 fold-change ≥ 0.58 were considered significantly deregulated. Gene lists derived from DOX and JQ1 coefficients were used for downstream analyses.

**Over-Representation Analysis (ORA)**

Pathway analysis was performed using over-representation analysis (ORA) with a per-pathway hypergeometric test. Gene sets were obtained from the Molecular Signatures Database (MSigDB) via the msigdbr R package (version 10.2.0), including hallmark (H), curated (C2), and ontology (C5) collections [9]. ORA was conducted using functions from the clusterProfiler R package [10]. Bonferroni-adjusted P-values ≤ 0.05 were considered statistically significant.

**ChIP-qPCR and ChIP-seq Procedure**

A total of $15 \times 10^6$ LN-2540GS and LN-2669GS spheroids were seeded in three 15-cm Petri dishes (TPP™, 93150) and cultured for 5 days. Fixation and chromatin shearing followed the "ChIP-IT® Express: Supplemental Protocol for Suspension Cells" using the ChIP-IT® Express Magnetic Chromatin Immunoprecipitation and Sonication Shearing Kit (Active Motif, 53008). Nuclei were isolated by lysis passing through a 29-gauge syringe (BD, 324827) 10 times and sheared in Bioruptor® Pico microtubes (Diagenode) using 6 (LN-2540GS) or 8 (LN-2669GS) cycles of 30 s on/off to obtain 100–600 bp fragments. Fragment size was verified by agarose gel electrophoresis, and chromatin was stored at –80 °C until use.

For ChIP-qPCR, 7 µg of chromatin per 200 µL reaction was used. Immunoprecipitation was performed with antibodies listed in Table S3, following the kit protocol. Purified DNA and input samples were quantified using a Qubit™ dsDNA HS Assay Kit. qPCR was performed with primers from Table S2. Percent enrichment over input was calculated using primer amplification efficiency, and fold enrichment over IgG was determined using the Active Motif protocol. Quality control was performed using the ChIP-IT® Control Human qPCR Kit (Active Motif, 53026).

ChIP-seq samples were prepared similarly, using 10 µg of chromatin per reaction and antibodies listed in Supplementary Table S3. Immunoprecipitated and input DNA were quantified using the Qubit™ dsDNA HS Assay Kit. A quality control ChIP-qPCR using the ChIP-IT® Control Human qPCR Kit was performed to verify enrichment prior to library preparation. ChIP libraries were prepared with the Ovation® Ultralow V2 DNA-Seq Library Preparation Kit (Tecan Life Sciences) by the GTF (University of Lausanne) and sequenced on an AVITI™ Sequencer as 150-nt paired-end reads, with 20 million reads for CTCF and Input samples, and 30 million reads for H3K27ac, H3K27me3, and H3K4me3 samples.

ChIP-seq data were processed using the nf-core/chipseq pipeline (v2.0.0) [11]. Reads were aligned to hg38 with Bowtie2 (v2.4.4), and peaks were called using MACS2 (v2.2.7.1). CTCF data were analyzed using the sevenC R package to compute correlation-based chromatin contacts, and to identify interactions within the *HOXA* cluster [12].

The regions of ChIP-Seq enrichment for the H3K27ac peaks outside of promoters (e.g. a region not contained within ± 2.5kb region flanking the gene promoter) are defined as active enhancers. The quantified peaks were merged into larger regions if they were less than 12.5 kb apart for the detection of the super-enhancers and the counts within the individual peaks were summed. These stitched regions were ranked by their counts and the threshold to classify super-enhancer and active enhancer was determined by ROSE algorithm [13].

#### ATAC-seq Sample and Library Preparation

ATAC-seq experiments were paired with our ChIP-seq experiments and performed with Active Motif's ATAC-seq Kit (Active Motif, 53150). Thus, $1\times10^6$ LN-2540GS and LN-2669GS spheroids were seeded in a 10cm petri dish and collected simultaneously. To remove potential artifacts caused by the potential presence of apoptotic cells, spheroids were incubated in 0.1mg/mL DNase I Solution (StemCell Technologies, 07900) with PBS at RT for 15min. Three aliquots of $1\times10^5$ cells from each sphere line were processed separately through the lysis step, and then pooled for tagmentation, PCR amplification, and library preparation. Next, DNA libraries were cleaned with Agencourt AMPure XP beads (Beckman Coulter, A63880) using a

1.8X bead ratio to remove adapter and primer dimers. Tagmentation and library preparation efficiencies were then tested with small aliquots on a Fragment Analyzer. ATAC libraries were sent to the GTF (University of Lausanne) and sequenced on an AVITI™ Sequencer as 150-nt paired-end reads at a depth of 100 million reads per sample.

The preprocessing of ATAC-seq data was performed following the pipeline suggested by nf-core/atacseq (version 2.1.2, https://nf-co.re/atacseq/2.1.2/) [11]. The sequencing data were aligned using Bowtie2 (version 2.4.4) with the NCBI hg38 genome reference assembly. Peak identification was performed using the peak caller Macs2 (version 2.2.7.1).

**Promoter Capture Micro-C**

LN-2540GS and LN-2669GS spheroids were cryopreserved and shipped to Cantata Bio for processing (CA, USA). Libraries were prepared using the Dovetail® Micro-C and Human Pan Promoter Panel Kits and sequenced to ~150 million paired-end reads. Reads were aligned to the GRCh38 genome reference (https://www.ncbi.nlm.nih.gov/datasets/genome/GCF_000001405.40). Contact matrices were imported and processed using the *cooler* Python package (v0.9.3) and R packages *HiCExperiment*, *HiContacts*, and *HiCcompare*. A/B compartments were predicted using constrained K-means in *HiCDOC* at 512kb resolution [14]. TADs were identified with *SpectralTAD* at 32kb and 64kb resolutions and compared using *TADCompare* [15,16]. The differential comparison between the LN-2540GS and LN-2669GS sphere lines was based on Promoter Capture Micro-C datasets and was normalized using a cyclic LOESS method at 8k resolution for each chromosome containing *HOX* genes. The interactions between two regions were tested by exact binomial tests for each pair. The differential comparison analyses were performed by functions from the multiHiCcompare R package (https://github.com/dozmorovlab/multiHiCcompare) [17].

### 2. Supplementary Tables

**Table S1.** List of primers used for RT-qPCR, InFusion molecular cloning, or ChIP-qPCR.

| Target | Method | Forward primer (5'-3') | Reverse Primer (5'-3') |
|---|---|---|---|
| *GAPDH* | RT-qPCR | AGGTGAAGGTCGGAGTCAACG | CGTTCTCAGCCTTGACGGTG |
| *HEXIM1* | RT-qPCR | AAGGACTAGCTAAAGGCGTCA | TGGCTAGTAGAGTCCTCGAAGTT T |
| *HOXA10* | RT-qPCR | AAGGTGAAAACGCAGCCAAC | CTAATCTCTAGGCGCCGCTC |
| *HOXA9* | RT-qPCR | TAAACCTGAACCGCTGTCGG | GCCTTCGCTGGGTTGTTTTT |
| *HOXA7* | RT-qPCR | CACCGAGCGCCAGATTAAGA | CCCCTCATTCCTCCTCGTCT |
| *HOXA6* | RT-qPCR | AAAGGCGGGCGAGTAGATG | CAGGCGGGGAGAAAAGTTGG |
| *HOXA5* | RT-qPCR | GTTACCTGACCCGCAGAAGG | CGCTCAGATACTCAGGGACG |
| *LAMINB1* | RT-qPCR | GTATGAAGAGGAGATTAACGAGAC | TACTCAATTTGACGCCCAG |
| pCW22-HOXA10-tGFP InFusion | *HOXA10-tGFP* cloning | ATCCTCTAGAGTCGACATGTCATG CTCGGAGAGCCC | CCAAGCTTGCATGCCTGCAGGT CTTTCTTCACCGGCATCTGC |
| Colony PCR (cloning validation) | *HOXA10-tGFP* cloning | CCCGCATCGAGAAGTACGAG | TCACCCAAGTCCCGTCCTAA |
| GAPDH-2 | ChIP-qPCR | undisclosed - Active Motif ChIP-IT® qPCR Analysis Kit | undisclosed - Active Motif ChIP-IT® qPCR Analysis Kit |
| Negative-1 | ChIP-qPCR | undisclosed - Active Motif ChIP-IT® qPCR Analysis Kit | undisclosed - Active Motif ChIP-IT® qPCR Analysis Kit |
| canonical *HOXA10* promoter | ChIP-qPCR | CCCGAGCTGATGAGCGAGTC | GCCAAATTATCCCACAACAATGT C |
| alternative *HOXA10* promoter 1 | ChIP-qPCR | TACCGTGCTGCTCTTAGT | TCCTCTCTCCTTCTCTCTCT |
| alternative *HOXA10* promoter 2 | ChIP-qPCR | AGAGAGAAGGAGAGAGGAGA | GGAATGCGCCGCTATAAA |
| alternative *HOXA10* promoter 3 | ChIP-qPCR | CGAATGCGCGTTGCTTTA | GCCTTTGTTCGCTTCTGATAC |

**Table S2.** List of shRNA constructs.

| Name | Clone ID | Manufacturer | Targeted mRNA sequence |
|---|---|---|---|
| shNTC | VSC11653 | Horizon Discovery Ltd | undisclosed |
| shHOXA10#2 | V3IHSHEG_491598 | Horizon Discovery Ltd | ACCTTACTCGAGAGCGGCG |
| shHOXA10#5 | V3IHSHEG_7259938 | Horizon Discovery Ltd | TGAATCGAGAAAACCGGAT |

**Table S3.** List of antibodies used for Western Blotting or Chromatin Immunoprecipitation.

| Antibody | Method | Reference | Manufacturer | Concentration |
|---|---|---|---|---|
| anti-GAPDH | Western Blot | G9545 | Sigma-Aldrich | 1:1'000 |
| anti-HOXA10 | Western Blot | #58891 | Cell Signaling Technology | 1:1'000 |
| anti-HOXA5 | Western Blot | ab140636 | Abcam | 1:1'000 |
| anti-Lamin B1 | Western Blot | #13435 | Cell Signaling Technology | 1:1'000 |
| anti-tGFP | Western Blot | TA150041 | OriGene | 1:1'000 |
| anti-H3 | Western Blot | #9715 | Cell Signaling Technology | 1:1'000 |
| anti-Rabbit IgG HRP | Western Blot | W4011 | Promega | 1:5'000 |
| Goat Anti-Mouse IgG HRP | Western Blot | 31430 | Invitrogen | 1:10'000 |
| anti-H3K9ac | ChIP | 91103 | Active Motif | 4µg/mL |
| anti-H3K27ac | ChIP | 91193 | Active Motif | 4µg/IP |
| anti-H3K27me3 | ChIP | 9733S | Active Motif | 1:50 |
| anti-H3K4me3 | ChIP | 91263 | Active Motif | 4µg/mL |
| anti-RNA Pol II + bridge | ChIP | 53026 | Active Motif | 2µg/IP |
| anti-IgG | ChIP | 53026 | Active Motif | 2µg/IP |
| anti-CTCF | ChIP | 91285 | Active Motif | 4µg/IP |

**Table S4.** Analysis of the association between *HOXA10* kockdown and JQ1 treatment and gene expression.

Analyses were based on a mixed generalized linear model (GLMM) with negative binomial distribution and random intercepts related to the cell types and the replicates.

See the separate Excel file for data.

**Table S5.** Over-representation analysis (ORA).

ORA for pathways associated with knock-down of *HOXA10* (DOX) or JQ1 treatment based on Hallmarks-C2 and C5 categories from the MsigDB database.

See the separate Excel file for data.

**Table S6.** Differential local genomic interactions for the LN-2540GS and LN-2669GS (Diff-HiC)

See the separate Excel file for data.

### 3. Supplementary Figures

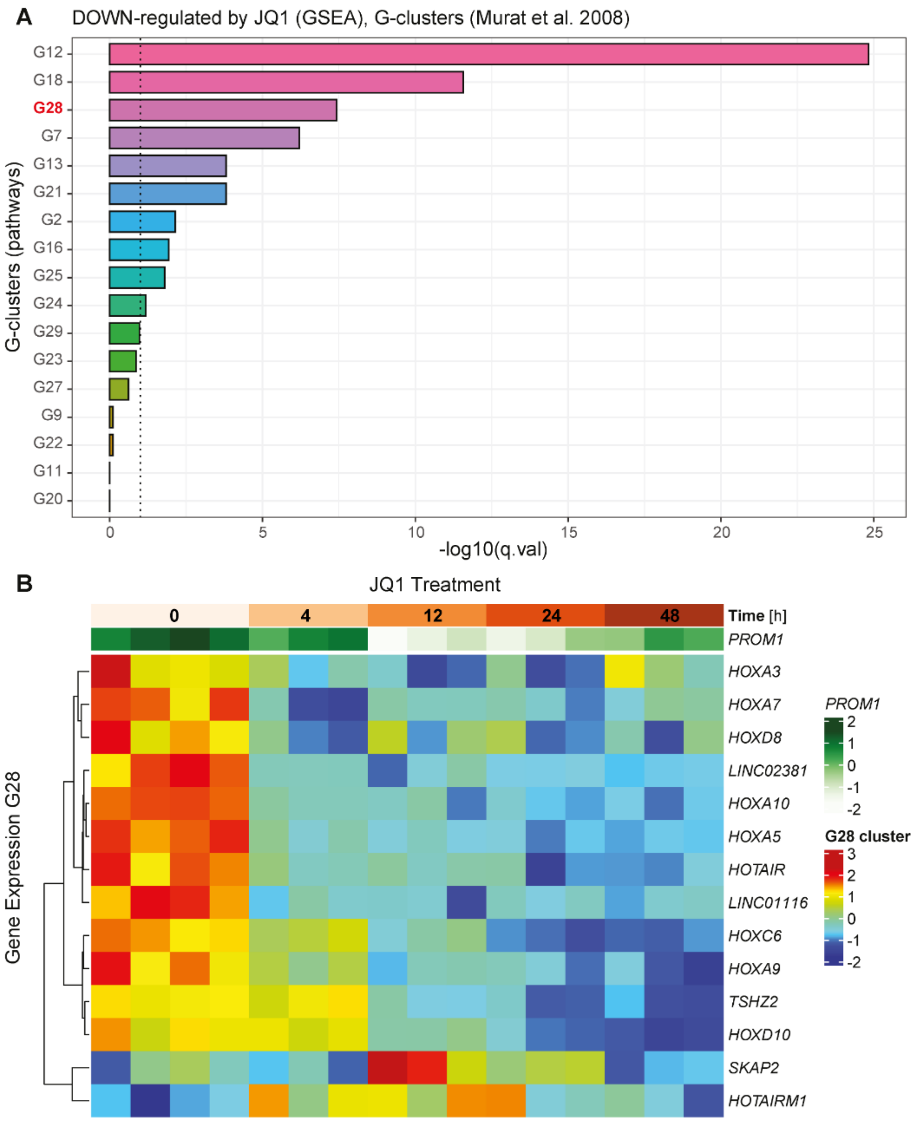

**Figure S1. BETi reveals coordinated repression of the *HOXA*-dominated, *HOX* signature in glioblastoma.**

The data set of differentially expressed genes extracted from the BETi-treated (1 µM JQ1) sphere line LN-2683GS [5,18] was subjected to Gene Set Enrichment Analysis (GSEA) using glioblastoma-derived gene sets from Murat *et al.* [19] (**A**). The gene set **G28** (marked in red), emerged as significantly enriched. The dotted line represents the cutoff for multiple testing correction. G28 corresponds to the hemopoietic stem-cell-related *HOX* signature that, when overexpressed, is associated with treatment resistance of GB [19]. (**B**) The heatmap visualizes the expression of the BETi-treated sphere line LN-2683GS over a 48 h time course, using the intersection of the G28 signature genes and the genes that passed the filter to enter the analyses. Coordinated repression of the *HOXA*-gene-dominated signature by JQ1 is observed. Expression of the stem cell marker *PROMININ 1* (*PROM1*) shows a similar profile under JQ1 treatment. *PROM1,* shown in green (not active in the clustering), is related to the signature G28 [19]. Of note, *SKAP2* and *HOTAIRM1* were not among the JQ1-differentially expressed genes (n=4712), as reflected in the heatmap.

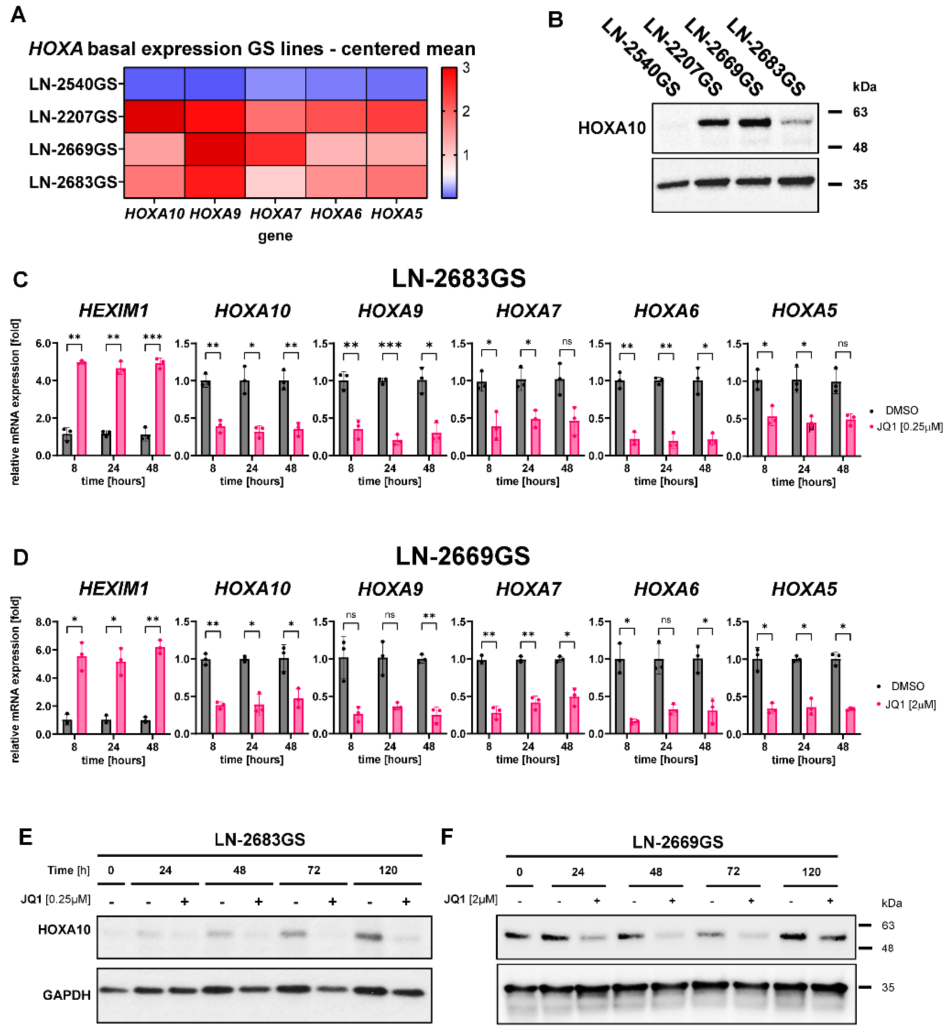

**Figure S2. BETi represses *HOXA* cluster gene transcription in high-*HOX* sphere lines LN-2683GS and LN-2669GS.**

**(A)** Baseline expression (qRT-PCR) of selected *HOXA* genes is illustrated in a heatmap for the GS-lines LN-2540GS, LN-2207GS, LN-2669GS, and LN-2683GS (normalized to mean *GAPDH* expression and mean-centered). **(B)** Corresponding baseline HOXA10 protein expression in the GS lines was determined by Western in relation to GAPDH. **(C, D)** The two high-*HOX* GS lines, LN-2683GS and LN-2669GS, were treated with JQ1 for 8 to 48 h at the concentrations indicated. Relative expression of selected *HOXA* genes was quantified by qRT-PCR, normalized to the mean of *GAPDH*, and the corresponding controls (DMSO) of each time point. Induction of *HEXIM1* expression by JQ1 served as a control for target engagement. Results are shown as mean values of biological replicates (n = 3). Error bars indicate SD. Two-way ANOVA was used to test treatment responses, and Šídák's multiple comparisons test for post hoc analysis. **(E, F)** HOXA10 protein expression was determined by Western for **(E)** LN-2683GS and **(F)** LN-2669GS over a time course of 120 hours of JQ1 treatment. The data shown are representative of biological replicates (n = 3). ns = non-significant, * = ($p \leq 0.05$), ** = ($p \leq 0.01$), *** = ($p \leq 0.001$).

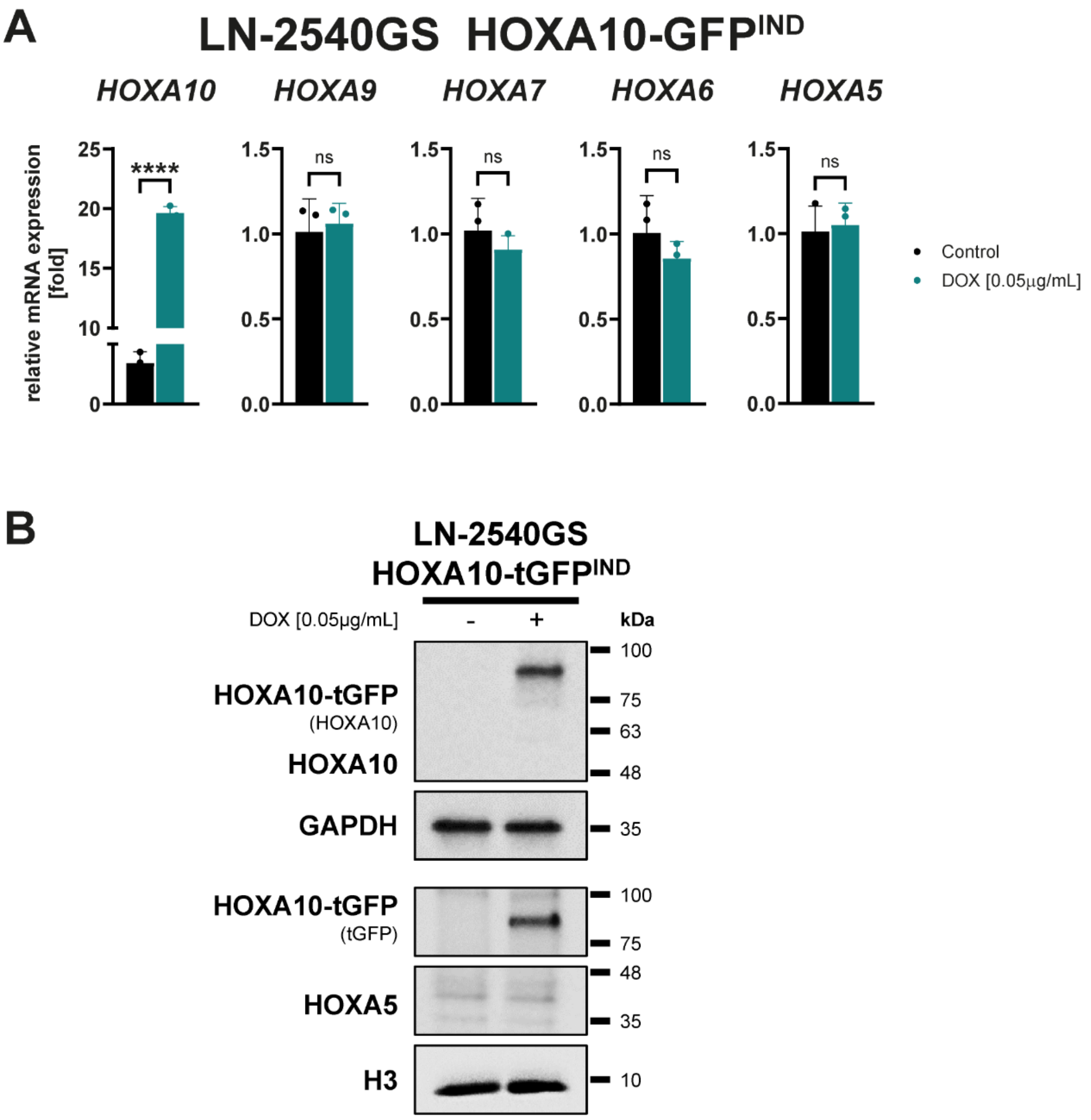

**Figure S3. Ectopic HOXA10 expression does not affect the expression of other *HOXA* genes in the low-*HOX* LN-2540GS.**

**(A)** Expression of selected *HOXA* genes was measured in the low-*HOX* LN-2540GS *HOXA10-tGFP*^IND spheres 48 h after induction of ectopic expression of *HOXA10* (0.05µg/mL doxycycline, DOX). The data were normalized to the mean *GAPDH* expression and the corresponding "Controls". Results are shown as mean values of biological replicates (n = 3). Error bars indicate SD. Treatment responses were tested by an unpaired t-test with Welch's correction. **(B)** Corresponding HOXA10-tGFP, endogenous HOXA10, and HOXA5 protein expression were determined by Western, controlled by GAPDH or HISTONE 3 (H3). The data shown are representative of biological replicates (n = 3). ns = non-significant, * = ($p \leq 0.05$), ** = ($p \leq 0.01$), *** = ($p \leq 0.001$), **** = ($p \leq 0.0001$).

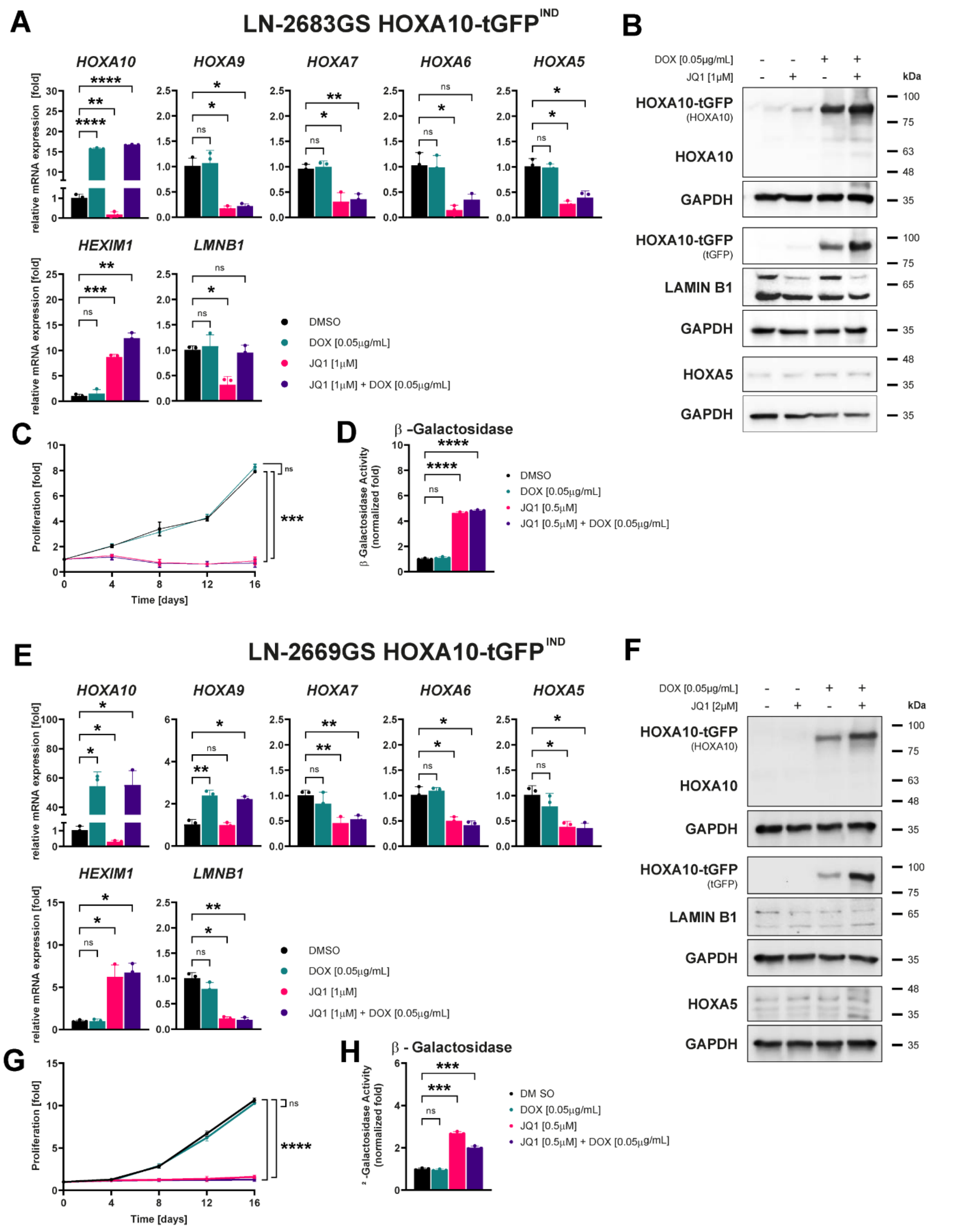

**Figure S4. Ectopic expression of HOXA10 in JQ1-treated high-*HOX* LN-2669GS neither restores expression of *HOXA* genes, nor proliferation, nor rescues cells from senescence.**

The high-*HOX* GS line LN-2669GS **(A)** transduced with inducible *HOXA10* (*HOXA10-tGFP*$^{IND}$) was pretreated for 72 hours with JQ1 at the concentrations indicated, followed by induction of ectopic expression of *HOXA10* for 48 h (doxycycline, +/-DOX). Expression of *HOXA10* and a selected set of *HOXA* genes, *HEXIM1,* and *LMNB1* (a measure of senescence) was analyzed by qRT-PCR. The data were normalized to the mean *GAPDH* expression and the corresponding "Control" values (DMSO). Results are shown as mean values of biological replicates (n = 3). Error bars indicate SD. Browne-Forsythe and Welch ANOVA tested treatment responses, and Dunnett's T3 multiple comparisons test was used for post hoc analysis. **(B)** Corresponding protein expression levels of ectopic and endogenous HOXA10, HOXA5, and LAMINB1 were determined by Western blot analysis, normalized to GAPDH, and a representative of biological replicates is shown (n = 3). **(C)** The relative proliferation rates of the high-*HOX* GS line LN-2669GS *HOXA10-tGFP*$^{IND}$, with ectopic expression of *HOXA10* (+/-DOX), were followed over 16 days of JQ1 treatment (as indicated) or DMSO. Fold changes are shown relative to day 0 as mean values of biological replicates (n = 3). Error bars indicate SD. Two-way ANOVA tested treatment responses on day 16, and Dunnett's multiple comparisons test was used for post hoc analysis. **(D)** To estimate senescence, relative β-Galactosidase activity was measured in the high-*HOX* sphere line treated as in **(C)**. Luminescence was measured on day 16, and absolute values were normalized to cell number, expressed as mean values from biological replicates (n = 3). Error bars indicate SD. Statistical analyses as in (A). **(E-H)** The same set of experiments as in **(A-D)** was performed in the high-*HOX* LN-2683GS *HOXA10-tGFP*$^{IND}$ spheres. ns = non-significant, * = ($p \leq 0.05$), ** = ($p \leq 0.01$), *** = ($p \leq 0.001$), **** = ($p \leq 0.0001$).

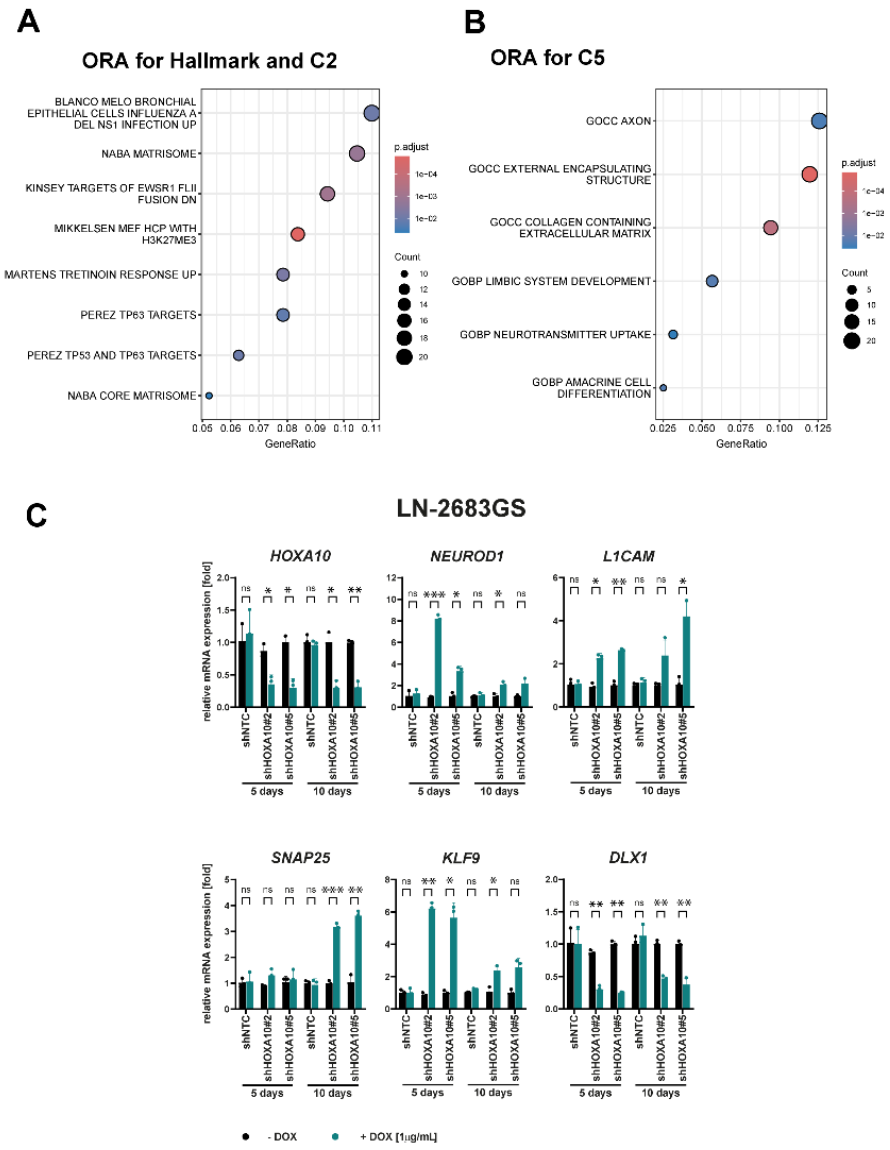

**Figure S5. *HOXA10* knockdown in high-*HOX* LN-2683GS affects stemness-, neurogenesis-, and extracellular matrix-associated signatures.**

Bubble plots showing significantly enriched gene sets 5 days after knock-down of *HOXA10* with the inducible (DOX+/-) sh*HOXA10*#2 and sh*HOXA10*#5 in LN-2683GS after overrepresentation analysis using **(A)** H and C2 or **(B)** C5 of the MSigDB collections. The color scale indicates the adjusted p-value (corrected for multiple comparisons using the Bonferroni method), with red indicating lower p-values, and the bubble size corresponding to the number of overlapping genes. **(C)** Validation of *HOXA10* knockdown and validation of expression changes in a selected set of candidate genes, as indicated, in LN-2683GS with inducible shNTC, sh*HOXA10*#2, or sh*HOXA10*#5 (induction for 5 or 10 days +/- DOX). The data were normalized to the mean *GAPDH* expression and the corresponding "Control" (-DOX values for each time point. Results are shown as mean values, with individual points representing separate biological replicates (n = 3). Error bars indicate SD. Treatment responses were tested by multiple unpaired t-tests with Welch's correction. ns = non-significant, * = ($p \leq 0.05$), ** = ($p \leq 0.01$), *** = ($p \leq 0.001$), **** = ($p \leq 0.0001$).

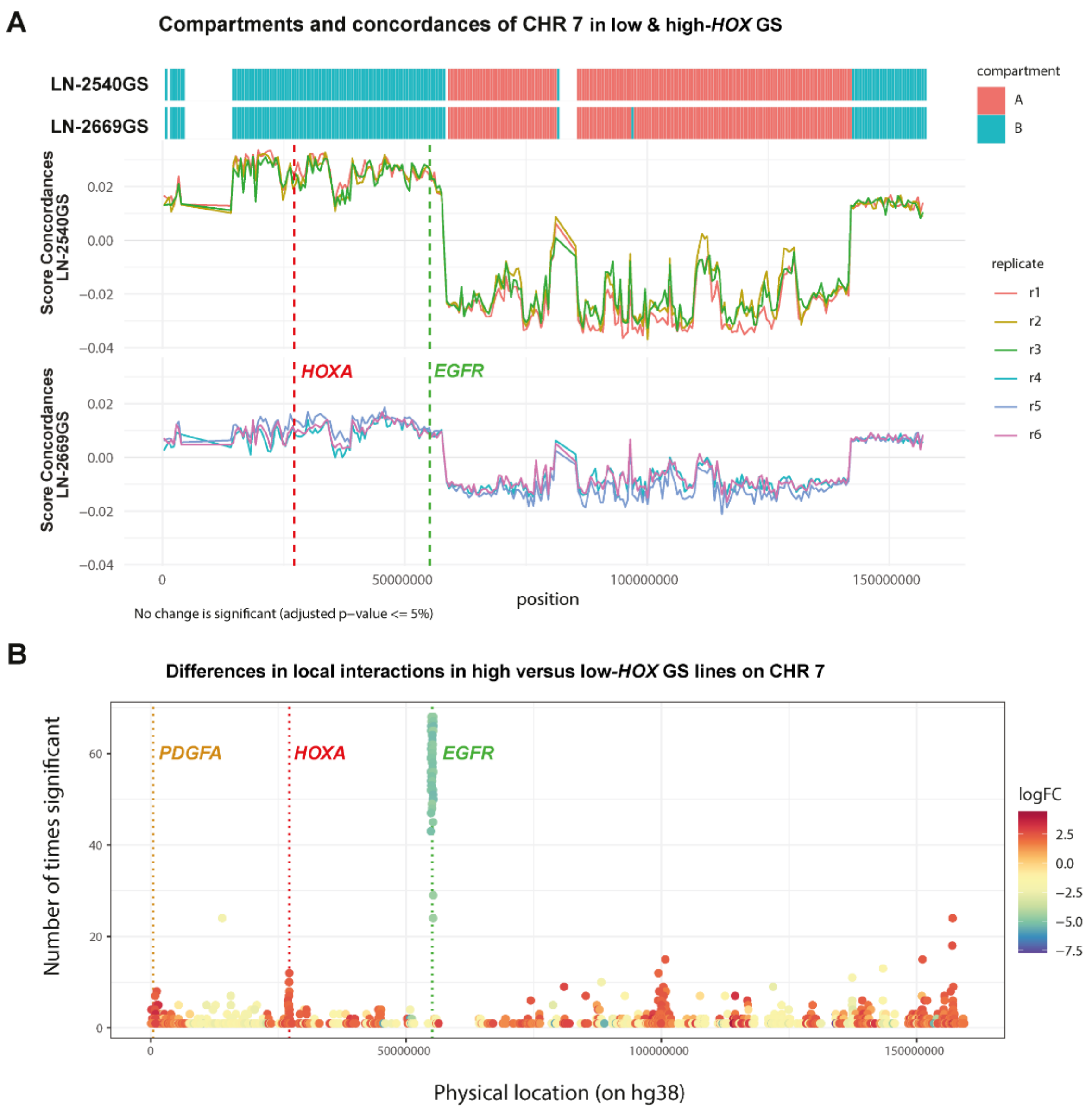

**Figure S6. Differential interaction analysis in the *HOXA* region in low- and high-*HOX* sphere lines.**

A/B compartment organization on Chromosome 7 (CHR 7) was interrogated by Hi-C. (**A**) A/B compartments on CHR 7 for the two sphere lines were described by concordance scores and classifications at a resolution of 512kb. The red and green dotted lines indicate the location of the *HOXA* cluster region and the *EGFR* gene, respectively. Of note, both GS lines show a gain of CHR 7 and a loss of CHR 10 (based on 450 k methylation data), hallmarks of GB. **(B)** Differences in local interactions in high versus low-*HOX* GS lines on CHR 7. Comparison of the interactions is represented as the number of times where the genomic location (1 anchor) is significant (absolute logFC > 1, logCPM >1, distance > 1, and Bonferroni's adjusted p-value < 0.1). The log fold change is visualized as a color gradient in the supplementary information. Values > 0 indicate increased local interactions in the high-*HOX* LN-2669G; values < 0 indicate increased interactions in the low-*HOX* LN-2540GS, respectively. The coordinates of the genomic locations are provided (GRCh38), and the loci of the *HOXA* cluster, *EGFR,* and *PDGFA* are annotated. Of note, the low-*HOX* GS line LN-2540GS has an *EGFR* amplification (usually organized extrachromosomal).

**Figure S7. Differential epigenetic landscapes at the *HOXC*, *D,* & *B* clusters of low and high *HOX* GS-lines.**

This Figure, related to Figure 5, compares the epigenetic landscapes in the *HOXC* (**A**), *HOXD* (**B**), *and HOXB* (**C**) regions between the low-*HOX* LN-2540GS and high-*HOX* LN-2669GS sphere lines.

**A**

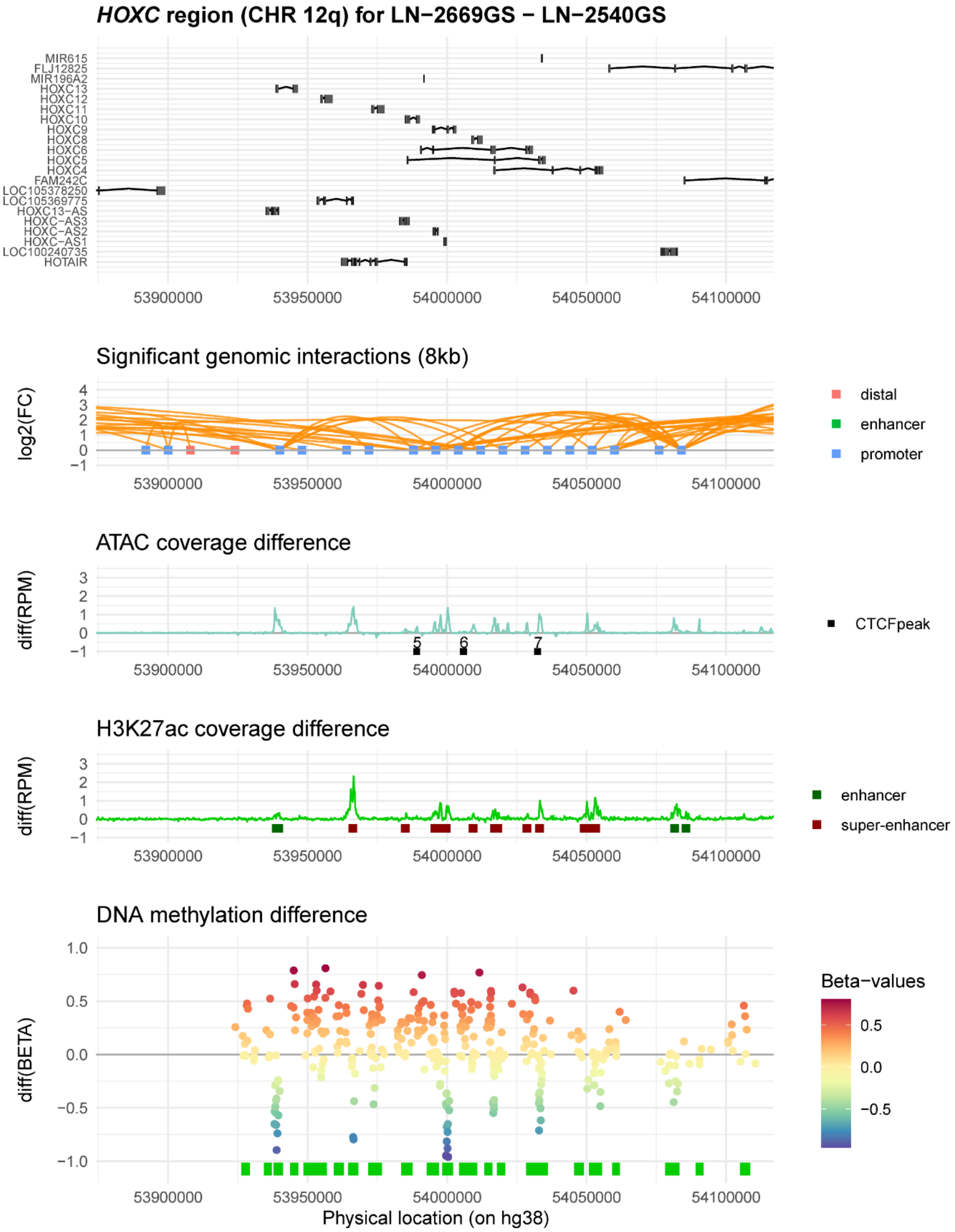

**A** Differential epigenetic landscape at the *HOXC* cluster of low-*HOX* LN-2540GS and high-*HOX* LN-2669GS sphere lines.

**B**

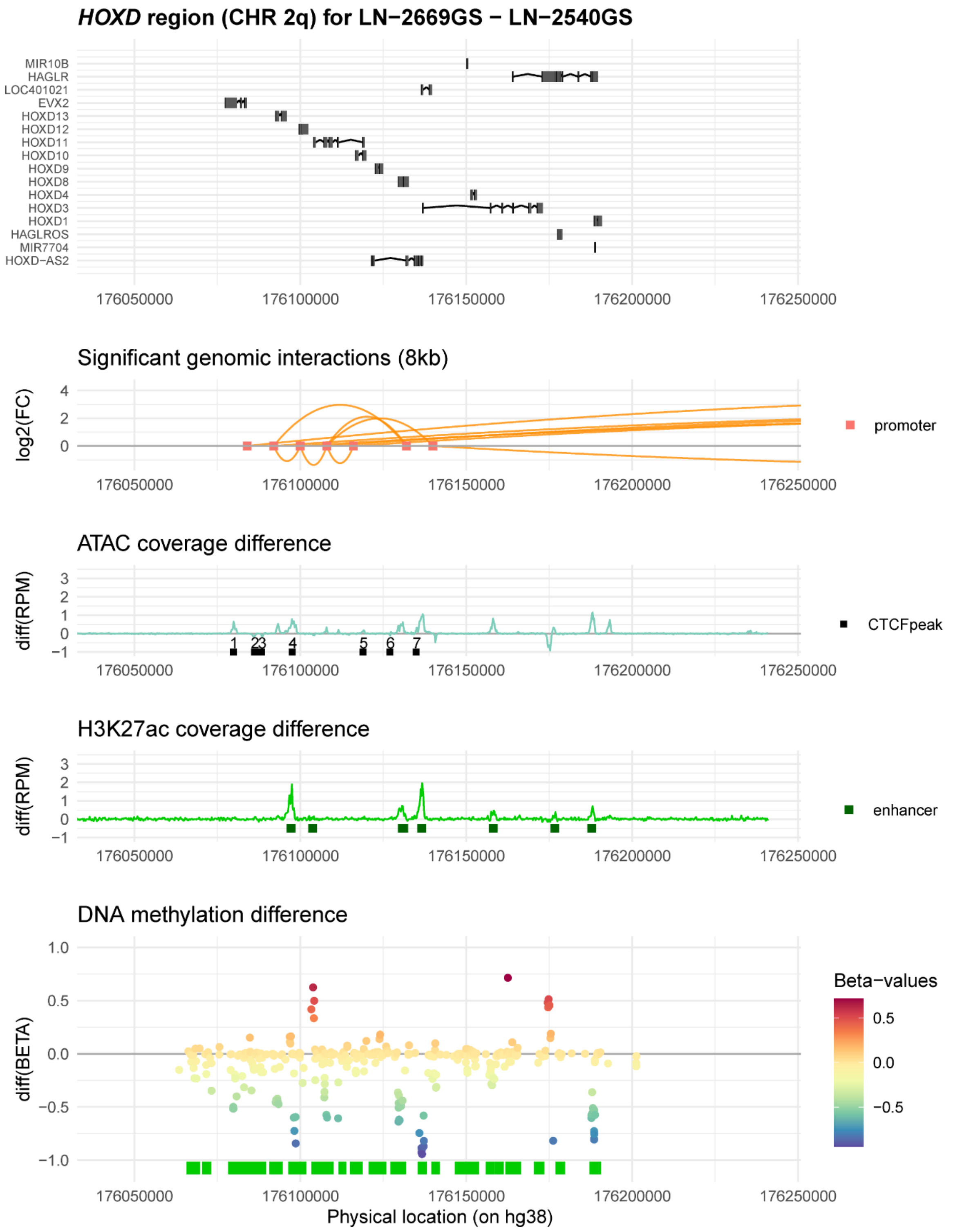

**B** Differential epigenetic landscape at the *HOXD* cluster of low-*HOX* LN-2540GS and high-*HOX* LN-2669GS sphere lines.

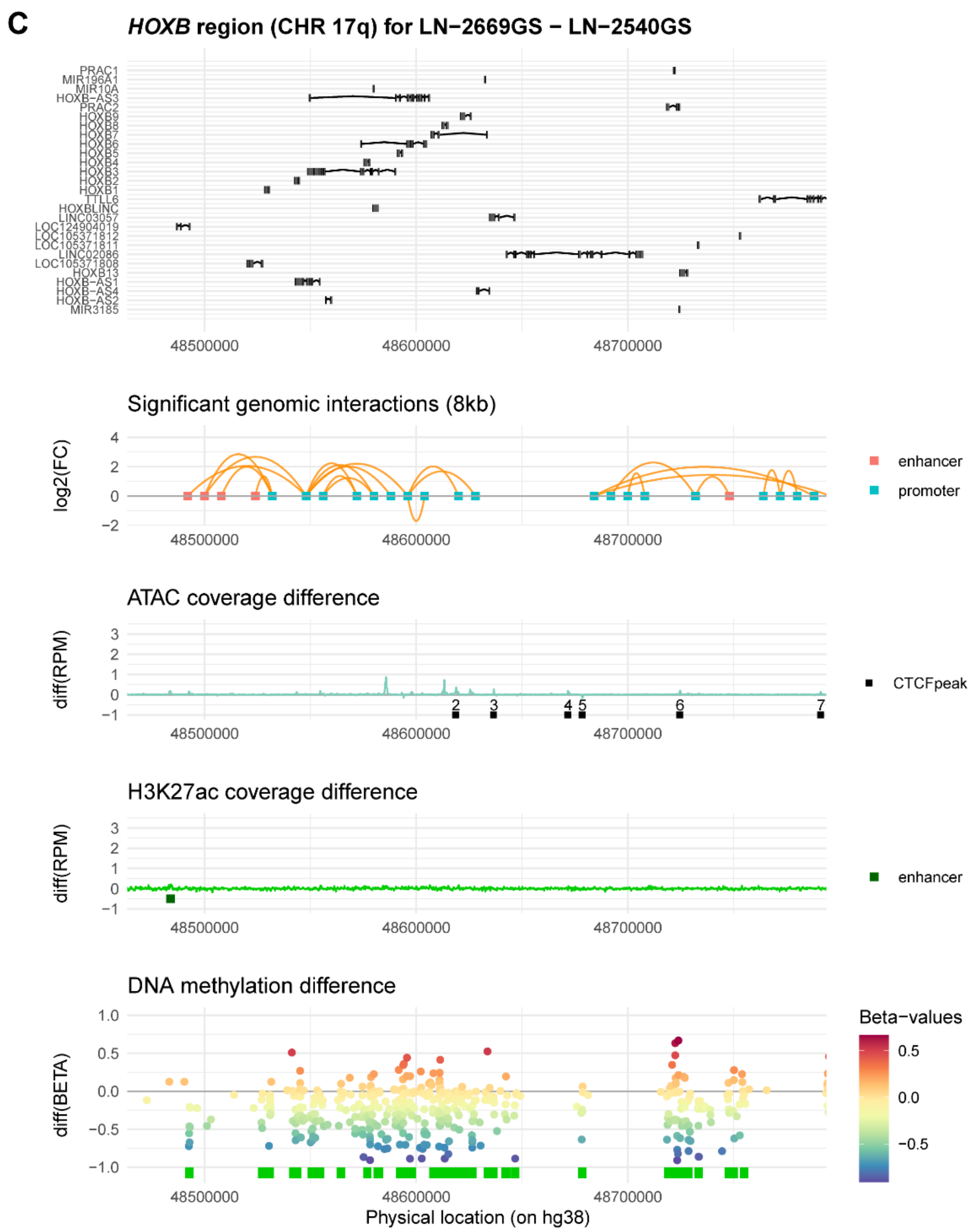

**C** Differential epigenetic landscape at the *HOXB* cluster of low-*HOX* LN-2540GS and high-*HOX* LN-2669GS sphere lines.

The epigenetic landscapes in the *HOXC* (**A**), *HOXD* (**B**), *and HOXB* (**C**) regions are compared between the low-*HOX* LN-2540GS and high-*HOX* LN-2669GS sphere lines. In brief, significant genomic interactions from the differential comparison (exact binomial test) between the two GS lines determined by micro-C are marked with orange arches. The summit corresponds to log fold-change (FC; values > 0, correspond to higher interactions in the high-*HOX* LN-2669GS). The annotation for each probe is indicated by a color code for promoter, enhancer, or distal. The difference of reads per million mapped reads (RPM) is provided for ATAC and H3K27ac coverage as a function of their physical location. The detected CTCF peaks and the enhancer classification by ROSE are superimposed on ATAC and H3K27ac tracks, respectively (enhancer, green; super-enhancer, red). The DNA methylation difference between the two sphere lines was computed with beta-values after functional normalization. CpG island locations are superimposed (green) on this track.

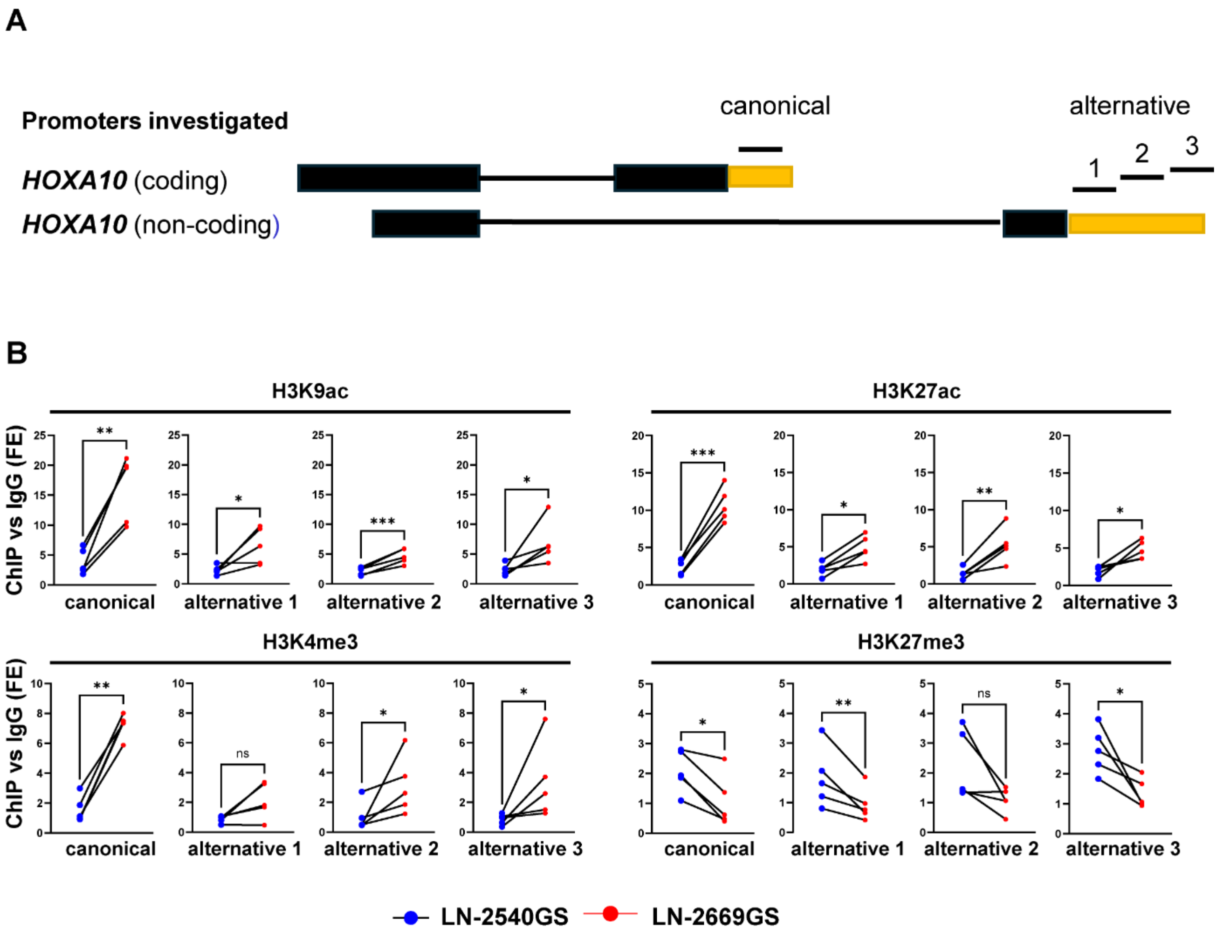

**Figure S8. Chromatin activation at the canonical and alternative *HOXA10* promoter in high and low-*HOX* GS-lines**

Chromatin marks were assessed at the canonical and alternative *HOXA10* promoter in low-*HOX* LN-2540GS and high-*HOX* LN-2669GS. **(A)** Graphic representation of the *HOXA10* promoter (canonical; alternative) regions targeted by ChIP-qPCR in **(B)**. **(B)** ChIP Fold Enrichment (FE) (normalized over IgG) tested by ChIP-qPCR with H3K9ac, H3K4me3, H3K27ac, and H3K27me3 marks in LN-2540GS (low-*HOX*) and LN-2669GS (high-*HOX*) spheres (n=5). Each point represents a biological replicate, with the lines connecting the paired biological replicates. A paired ratio t-test was conducted to test differences within each replicate. ns = non-significant, * = ($p < 0.05$), ** = ($p < 0.01$), *** = ($p < 0.001$).
